# SpiderLearner: An ensemble approach to Gaussian graphical model estimation

**DOI:** 10.1101/2021.07.13.452248

**Authors:** Katherine H. Shutta, Laura B. Balzer, Denise M. Scholtens, Raji Balasubramanian

## Abstract

Multivariate biological data are often modeled using networks in which nodes represent a biological variable (e.g., genes) and edges represent associations (e.g., coexpression). A Gaussian graphical model (GGM), or partial correlation network, is an undirected graphical model in which a weighted edge between two nodes represents the magnitude of their partial correlation, and the absence of an edge indicates zero partial correlation. A GGM provides a roadmap of direct dependencies between variables, providing a valuable systems-level perspective. Many methods exist for estimating GGMs; estimated GGMs are typically highly sensitive to choice of method, posing an outstanding statistical challenge. We address this challenge by developing SpiderLearner, a tool that combines a range of candidate GGM estimation methods to construct an ensemble estimate as a weighted average of results from each candidate. In simulation studies, SpiderLearner performs better than or comparably to the best of the candidate methods. We apply SpiderLearner to estimate a GGM for gene expression in a publicly available dataset of 260 ovarian cancer patients. Using the community structure of the GGM, we develop a network-based risk score which we validate in six independent datasets. The risk score requires only seven genes, each of which has important biological function. Our method is flexible, extensible, and has demonstrated potential to identify *de novo* biomarkers for complex diseases. An open-source implementation of our method is available at https://github.com/katehoffshutta/SpiderLearner.

## Introduction

Gaussian graphical models (GGMs) are a modeling framework for network-based analyses of multivariate data. This framework begins with the assumption that data are sampled from a multivariate normal distribution; under this assumption, a GGM is defined as a graph in which nodes correspond to variables and weighted edges correspond to the magnitude of the partial correlation between them (1). In this framework, the absence of an edge between nodes corresponds to zero partial correlation, i.e., conditional independence between the variables, given the other variables in the network.

The weighted adjacency matrix of a GGM consists of the partial correlations between nodes. Let **X** ~ *N_p_*(**0**, ∑) be a centered *p*-dimensional multivariate normal random variable, and let **X**_−*i*, −*j*_ represent **X** with the *i^th^* and *j^th^* variables removed. The partial correlation between *X_i_* and *X_j_* is defined as:

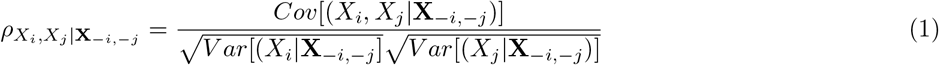

Under the assumption of multivariate normality, a particularly useful relationship holds between the precision matrix Θ = ∑^−1^ and the partial correlation (2). Let *θ_ij_* represent the *i, j^th^* element of Θ; it can be shown that

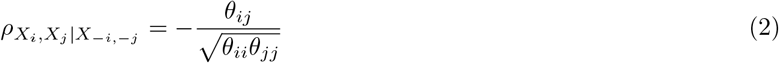

Equation 2 shows that estimating a GGM is equivalent to estimating Θ. In the case where the sample size *n* is much larger than the number of predictors *p*, a maximum likelihood estimate of Θ can be found simply by inverting the sample covariance. When *n* is close to or less than *p*, this inverse is undefined or numerically unstable, meaning this method cannot be used. The usual approach in this setting is the graphical lasso, which estimates a sparse precision matrix by optimizing the penalized likelihood function (3; 4; 5)

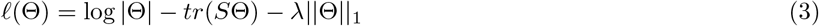

where *S* is the sample covariance matrix of the observed data and λ > 0 is a non-negative tuning parameter, with higher values of λ leading to sparser estimates of Θ. Several existing open-source software resources implement various versions and extensions of the graphical lasso, including methods for selecting the tuning parameter λ. For example, the `glasso` R package implements the original algorithm developed in 2008 by Friedman et. al. as augmented by computational advances developed in 2011 by Witten et. al. (3; 6). The `huge` R package incorporates the graphical lasso algorithm and additionally provides options for a tuning-insensitive method called `tiger` published by Liu et al. in 2017 (7),(8). The `bootnet` R package includes a broad range of different network estimation methods, seven of which are for GGM estimation, in a framework for bootstrap estimation of network accuracy (9).

There is clearly no shortage of options for a researcher who is interested in estimating a GGM; this is both a blessing and a curse. Estimating a GGM using these packages requires the researcher to make several decisions with regard to data preprocessing, tuning parameter selection, choice of scoring criteria for model selection, and selection of hyperparameters for these scoring criteria. The final estimated GGM may be highly sensitive to these choice, making it difficult to compare GGMs across studies and assess reproducibility (10; 11; 12). Because it is impossible to know *a priori* which approach is best for a given problem, researcher bias toward use of a particular “favorite method” can have a large impact on the estimation and interpretation of a GGM, consequently affecting the scientific conclusions inferred.

Ensemble methods are a broad class of statistical approaches which follow the general principle of combining several different candidate models to generate a single ensemble model (13; 14). One such method is the Super Learner approach of van der Laan et al (15). Super Learner uses an internal cross-validation scheme to fit a convex combination of candidate algorithms (“learners”) that minimizes a user-defined loss function. This convex combination is the Super Learner ensemble model. Large-sample properties of the Super Learner are established by comparison to the expected loss (i.e., risk) of an oracle model, which is the best model among all possible convex combinations given the true, unknown, data generating process. Under mild conditions on the loss function and the set of candidate learners, the expected difference between the risk of the Super Learner ensemble model and the risk of the oracle model converges to zero as the sample size goes to infinity(15).

Here, we develop SpiderLearner, a network estimation tool which applies the Super Learner approach to the problem of fitting a GGM by optimizing a likelihood-based loss function through the use of cross-validation. Our approach improves GGM estimation by circumventing the complicated decision-making burden described above. The SpiderLearner considers a library of candidate GGM estimation methods and constructs the optimal convex combination of their results, eliminating the need for the researcher to make arbitrary decisions in the estimation process. Through simulation studies, we demonstrate that the SpiderLearner achieves equal or better performance than each of the candidate approaches according to several criteria (out-of-sample likelihood, bias, mean squared error (MSE), matrix correlation).

Previous work has shown that the use of network models in prediction problems for disease outcomes is a promising area of work (e.g.,(16; 17; 18)). We connect this previous work to ours by presenting an illustrative application of the SpiderLearner to develop a robust risk score for ovarian cancer prognosis based on a publicly-available gene expression dataset (19; 20).

The SpiderLearner improves the applicability of GGMs as a network modeling framework, creating a more robust methodology by using data-driven ensemble learning to eliminate the need for researchers to choose an estimation approach. Our risk score application demonstrates the potential of the SpiderLearner to discover meaningful biological insights in complex multivariate data.

## Materials and methods

### SpiderLearner model formulation

The foundations for a Super Learner-type method are (i) specifying a library of candidate algorithms, (ii) specifying a loss function, and (iii) implementing a cross-validation scheme to determine the optimal convex combination of the candidates (21). We introduce the foundations of our method similarly, but focus first on (ii) and (iii); we address (i) when describing our simulation study design.

To develop the loss function for the SpiderLearner, we begin by supposing that we have a library of *M* different candidate methods and have applied each candidate to obtain *M* estimated GGMs for a given input dataset. Our next goal is to estimate a weighted combination of these *M* estimates that may provide an even better fit than each method does alone, in the spirit of the Super Learner approach(15). We consider first a basic test-train setting, in which we assume the availability of two independent datasets *X_train_* and *X_test_*, drawn at random from the same population. Let 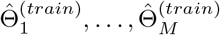 denote the estimated precision matrices from the application of the *M* different methods to *X_train_*. We seek a convex combination

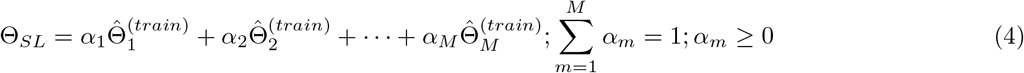

that minimizes the negative log-likelihood in the independent dataset *X_test_*:

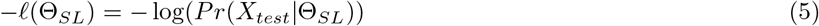

Let 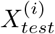 denote the data vector for the *i^th^* observation in the test dataset. In the GGM setting, where 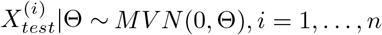, the log likelihood can be expressed as:

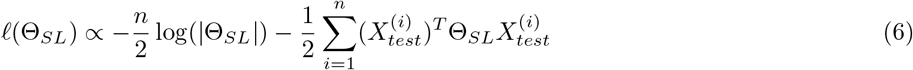

We negate the log likelihood and incorporate our definition of Θ*_SL_* from Equation 4 to develop a loss function in terms of the coefficients ***α*** = *α*_1_, …, *α_M_*:

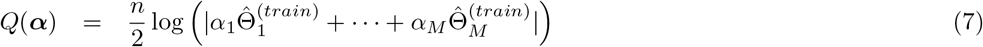

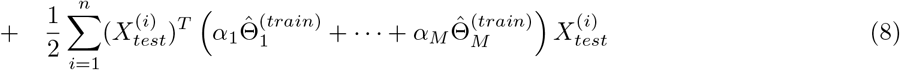

This loss function 12 is then minimized, subject to the constraints of the convex combination:

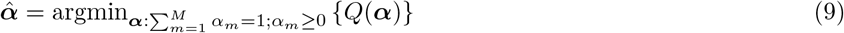

Standard constrained optimization algorithms can such as those implemented in the the `solnp` function from the R package `Rsolnp` (22) can be used to find the coefficients *α* that solve 9 on the test dataset. Once these coefficients have been found we complete the process by running the original *M* candidate methods again using the full dataset *X* = *X_train_* ∪ *X_test_*, obtaining estimates 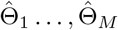. We then use these estimates to construct the SpiderLearner estimate of the precision matrix as:

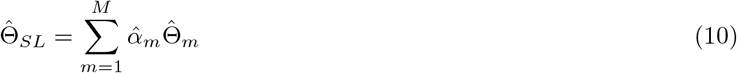

The train-test approach is limited in that the estimate of the out-of-sample loss will tend to be (i) an overestimate due to the relatively small size of the training dataset, and (ii) suffer from high variability due to the sensitivity of the approach to the characteristics of the training and test datasets (23). To overcome these limitations, we extend our approach from the simple train-test setting described above to *K*-fold cross-validation, where *K* > 2. *K*-fold cross-validation has the advantage of permitting the user to navigate the bias-variance tradeoff in the estimation of out-of-sample loss (23).

Briefly, we begin the *K*-fold cross-validation by partitioning the data *X* into *K* folds of approximately equal size ~ *n/K*. We next repeat the above process of determining the precision matrix estimator 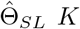 times; each time, data from the *k^th^* fold is withheld (*k* = 1,…, *K*) as the test set while the remaining (*K* – 1) of the folds serve as the training set.

To provide further detail, we first introduce some notation. Let *X_k_* be the *k^th^* fold of the dataset *X*, and let *X_−k_* be the remainder of the dataset *X* with the *k^th^* fold withheld. Let 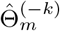 be the precision matrix estimate for method *m* trained on *X_−k_*, and let 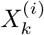 be the *i^th^* observation in fold *K*. We define 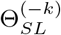, which is a function of ***α***, as:

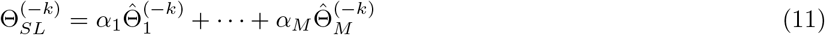

Next, we define *Q_k_*(***α***) to be the loss of the estimate 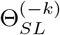 evaluated on the withheld data *X_k_*:

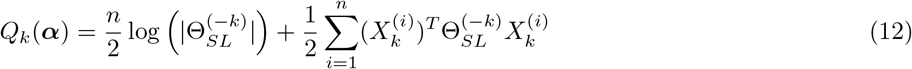

Let 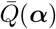 be the average loss aross *K* folds:

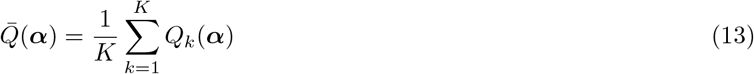

Let *n_k_* be the number of observations in the *k^th^* fold. Then the *K*-fold cross-validated coefficient estimator of 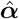 is:

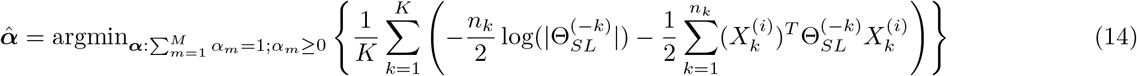

Finally, let 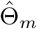 be the precision matrix estimate from method *m* using the full dataset. We then define the *K*-fold cross-validated SpiderLearner estimator as:

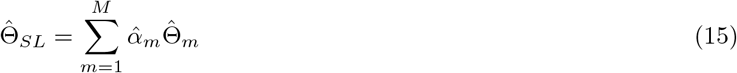

A diagram of this workflow for *M* = 4 estimation methods and *K* = 5 cross-validation folds is shown in Figure 1. The choice of *K* may depend on a variety of factors including sample size and number of predictors (i.e., dimensionality of the problem); in practice, *K* = 5 and *K* = 10 have demonstrated generally good balance in the bias-variance trade off(23). We discuss the choice of *K* further in the Results section.

**Figure 1:**
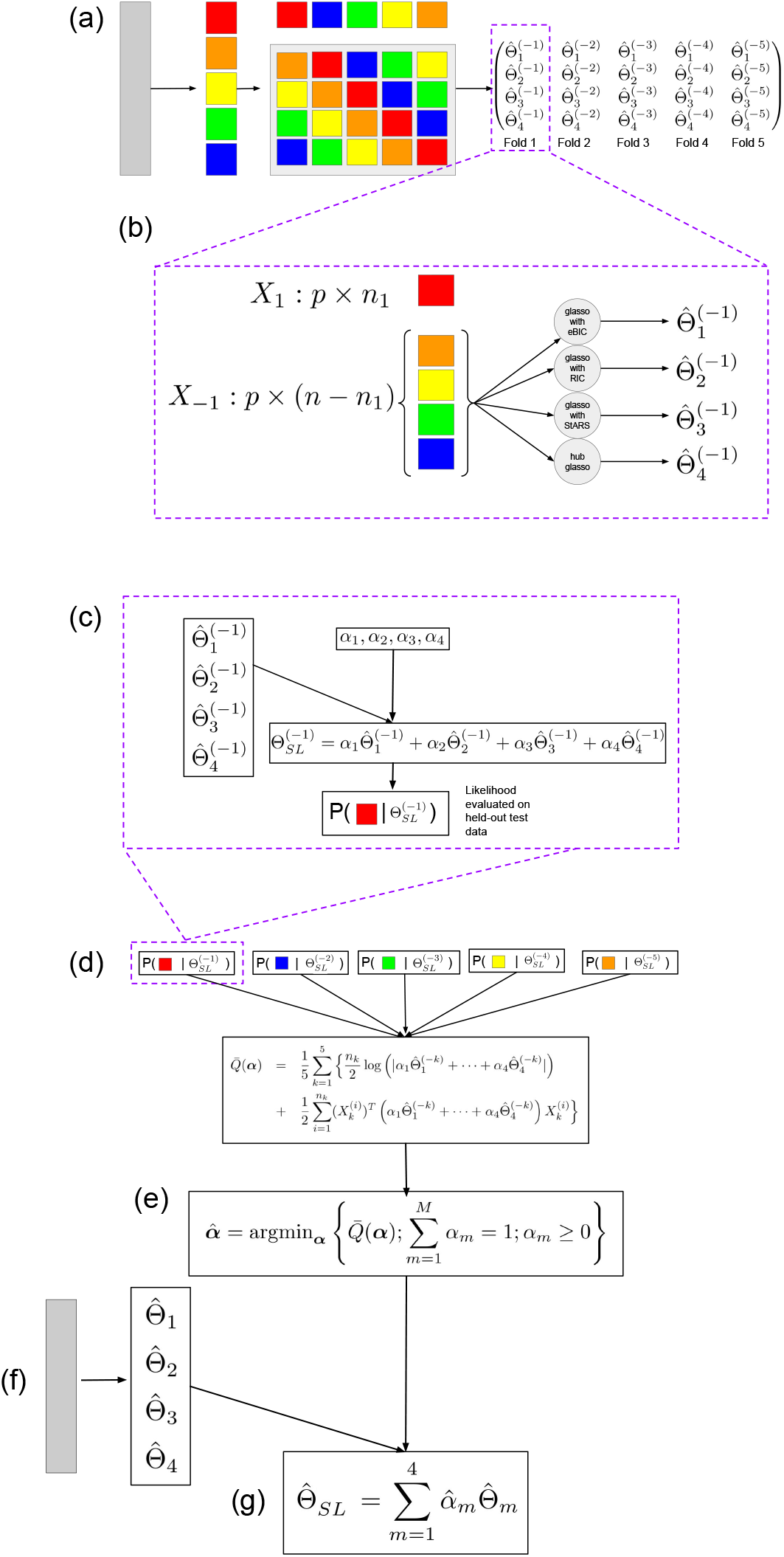
(a) Data are partitioned into five folds. Each fold is left out from the model fitting process in turn. (b) Every candidate model is fit on the training data in each fold. This generates an (*M* = 4) × (*K* = 5) array of estimated matrices. (c) For each held-out dataset *k* and coefficient set ***α*** = (*α*_1_,…, *α*_4_), the estimator 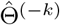 is calculated from the estimates obtained in (b). The likelihood of the estimator given the held-out data is then calculated. The process is repeated across all *K* = 5 folds and averaged to yield our loss function. (d) The loss function is minimized to yield the optimal coefficients 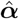, subject to the constraints of the convex combination. (e) The *M* = 4 methods are used to fit 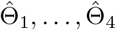 on the whole dataset. (f) The final SpiderLearner estimator 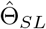 is calculated as the convex combination of the coefficients selected in (d) with the models fit in (e).

#### Large-sample properties

The large-sample properties of Super Learner derived by (15) require a bounded loss function. Our loss function 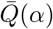 (Equation 12) is not bounded; therefore, it is not clear if the large-sample oracle results of (15) apply with the log likelihood-based loss function evaluated on multivariate normal data. (24) note that oracle results also hold for certain types of unbounded loss functions as described in (25); however, it is not straightforward to formally show that 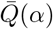 meets the necessary criteria (see S1 Appendix in the Supplement for details). We note this as an area for future work, while observing that in practice the log likelihood is often used as a loss function for Super Learner estimation (e.g., (26; 27; 28)), and the log likelihood loss is provided as part of the standard implementation of SuperLearner (29).

We additionally explored a transformation that permits the application of the oracle results of (15). Specifically, 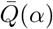 can be transformed into a bounded loss function 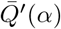 by applying the inverse logit function:

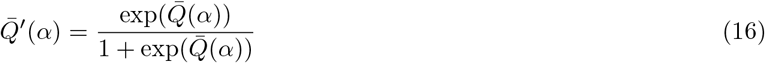

In practice, this transformation can be sensitive to the scale of 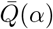 and quickly become numerically equal to one for a broad range of *α*. We observed good behavior by scaling 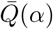 to the sample size *n* and number of predictors *p*:

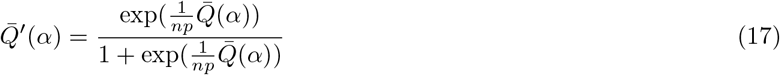

However, we suspect that further issues with the numerical stability of this transformation are possible and encourage diagnostics such as testing the value of 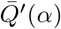 for different values of *α* when using this transformation in practice.

Equation 12 is used as the loss function in the simulation and application sections below, while the original loss function (Equation 12) and the bounded loss function (Equation 17) are both provided as options in the SpiderLearner implementation. A comparison of performance of the original and bounded loss functions can be found in Supplementary Figure S1.

### Simulation study

To assess the performance of the SpiderLearner algorithm, we conducted several simulation studies with varying sample sizes and numbers of predictors (Table 1a). In all simulations, a variety of network topologies and densities were considered (Table 1b). The simulation workflow (Figure 2) consisted of (i) designing gold-standard networks corresponding to each topology and density, (ii) assigning edge weights to the network based on an observed distribution of partial correlations from a real biological dataset and converting the associated weighted adjacency matrices to valid precision matrices, (iii) sampling multivariate normal data based on the precision matrices from (ii), (iv) using various methods, including our proposed ensemble method, to estimate the original network from the sampled data, and (v) comparing the estimated network to the original gold standard used to generate the data.

**Table 1:** Details of the simulation study designed to test the robustness of the SpiderLearner algorithm to differences in dimensionality and topology.

| Simulation | $n$ | $p$ | $q$ | $.9 * n/q$ |
| --- | --- | --- | --- | --- |
| A | 10,000 | 50 | 1275 | 7.06 |
| B | 1,600 | 50 | 1275 | 1.13 |
| C | 100 | 50 | 1275 | 0.07 |
| D | 60 | 100 | 5050 | 0.01 |

| Topology | Density | igraph Function | Simulated Density (A,B,C) | Simulated Density (D) |
| --- | --- | --- | --- | --- |
| Random | Low | `sample_gnp` | 0.053 | 0.061 |
| Random | High | `sample_gnp` | 0.219 | 0.194 |
| Small World | Low | `sample_smallworld` | 0.082 | 0.061 |
| Small World | High | `sample_smallworld` | 0.204 | 0.202 |
| Scale-Free | Low | `sample_pa` | 0.079 | 0.059 |
| Scale-Free | High | `sample_pa` | 0.192 | 0.191 |
| Hub-and-Spoke | Low | `sample_pa` | 0.079 | 0.059 |
| Hub-and-Spoke | High | `sample_pa` | 0.192 | 0.191 |

(b) Gold-standard networks were constructed using a variety of functions from the **igraph** package. Graph density is a function of the parameters used in each function as well as the number of predictors in the graph, and cannot be exactly specified. Parameters used in this study were chosen to achieve approximately 6 percent dense graphs in the low-density cases and 20 percent dense graphs in the high-density cases.

**Figure 2:**
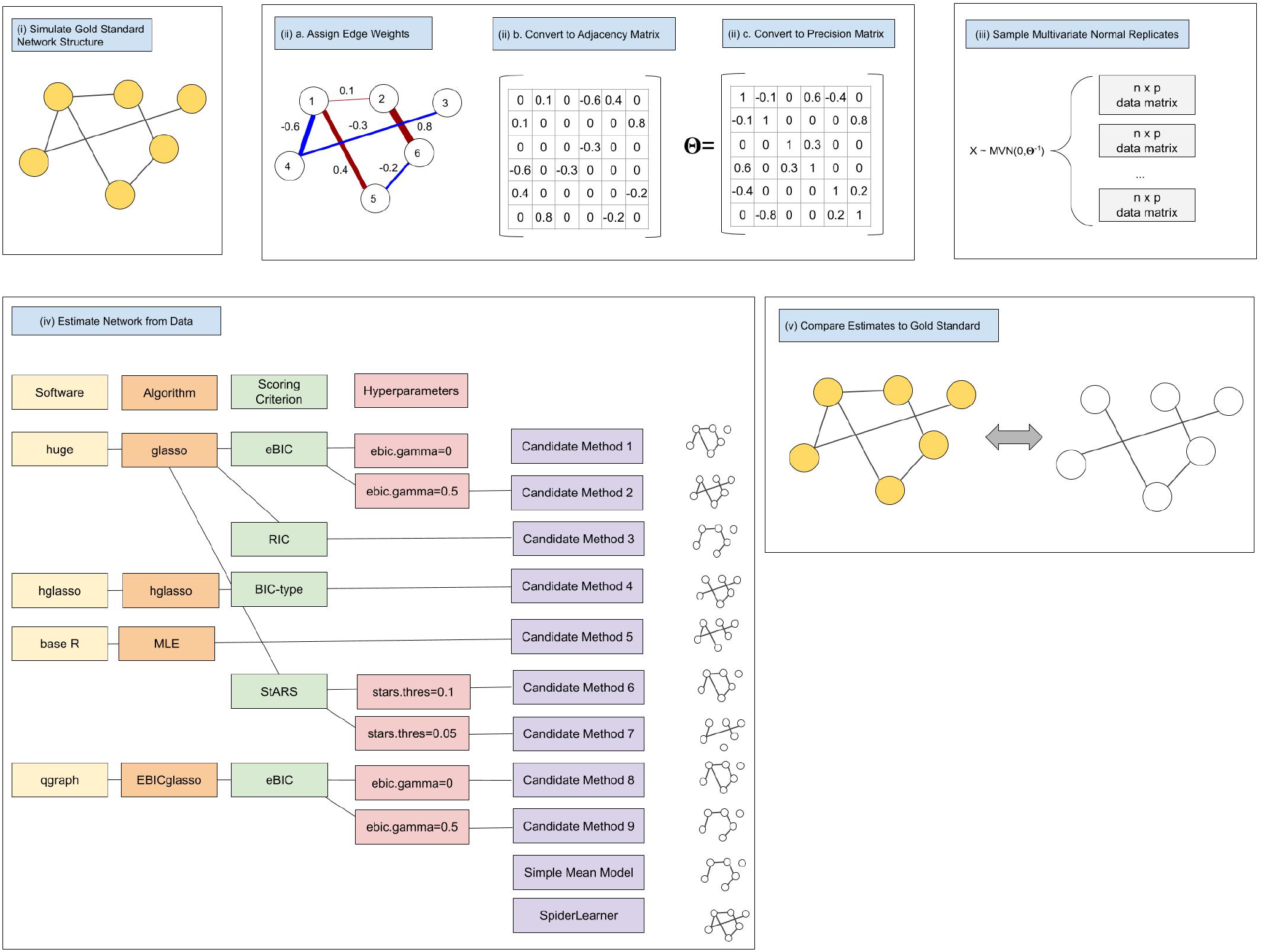
Simulation study workflow. In (i) we design gold-standard networks. In (ii), we assign edge weights to the gold standards by sampling from the distribution of partial correlations observed in the CATHGEN dataset and convert the corresponding adjacency matrices to precision matrices. In (iii), we sample multivariate normal data based on the precision matrices from (ii). In (iv), we estimate the networks from the sampled data. In (v), we compare the estimated network to the gold standard.

We explored four different network topologies in our simulations: random, small world, scale-free, and hub-and-spoke. Each topology has unique characteristics that may be relevant for biological data.

#### Random graph

In an Erdös-Renyí/Gilbert random graph on *p* nodes, it is assumed that each of the 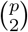 possible edges is equally likely to exist, according to some fixed probability π (30; 31). A random graph can be constructed by sampling each edge independently from a *Bernoulli*(π) distribution (31). The degree distribution of a random graph is approximately a Poisson distribution (32).

#### Small world graph

A small world graph is characterized by a type of community structure that is absent in the random graph and which results in a shorter average path length (33). A small world graph on *p* nodes can be simulated by beginning with a circular lattice in which each node is connected to *k* of its neighbors. From this point, the graph is “rewired” by going around the lattice and reconnecting each edge to a different node at random with some fixed rewiring probability π. These random disruptions of the lattice structure create shortcuts across the graph, leading to the lower average path length for the graph as a whole.

#### Scale-free graph

A scale-free graph is characterized by a power law degree distribution that can be simulated by considering an empty graph on *p* nodes and adding edges *k*-at-a-time following a growth and preferential attachment model, in which the probability that a particular node gets another edge added to it is proportional to how many edges it already has (34; 32). A log-log plot of the degree distribution (i.e., log frequency vs. log degree) in a scale-free graph is approximately a straight line; thus the model for generating this graph is referred to as linear preferential attachment.

#### Hub-and-spoke graph

A hub-and-spoke graph arises in a similar way as the scale-free graph does, but the probability that an edge is added to a particular node is proportional to the *k^th^* power of the degree of that node, for some *k* > 1 (superlinear preferential attachment) (32). This graph is characterized by hub nodes with very high degree and non-hub nodes with very low degree.

We explore these topologies along with graphs of different edge densities, where the edge density of a graph is defined as the number of edges divided by 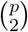, the number of possible edges on *p* nodes. For each of the four topologies, we simulated networks with two different density levels (low density: approximately 6 percent dense, and high density: approximately 20 percent dense). A visualization of these gold-standard networks shown in Figure 3.

**Figure 3:**
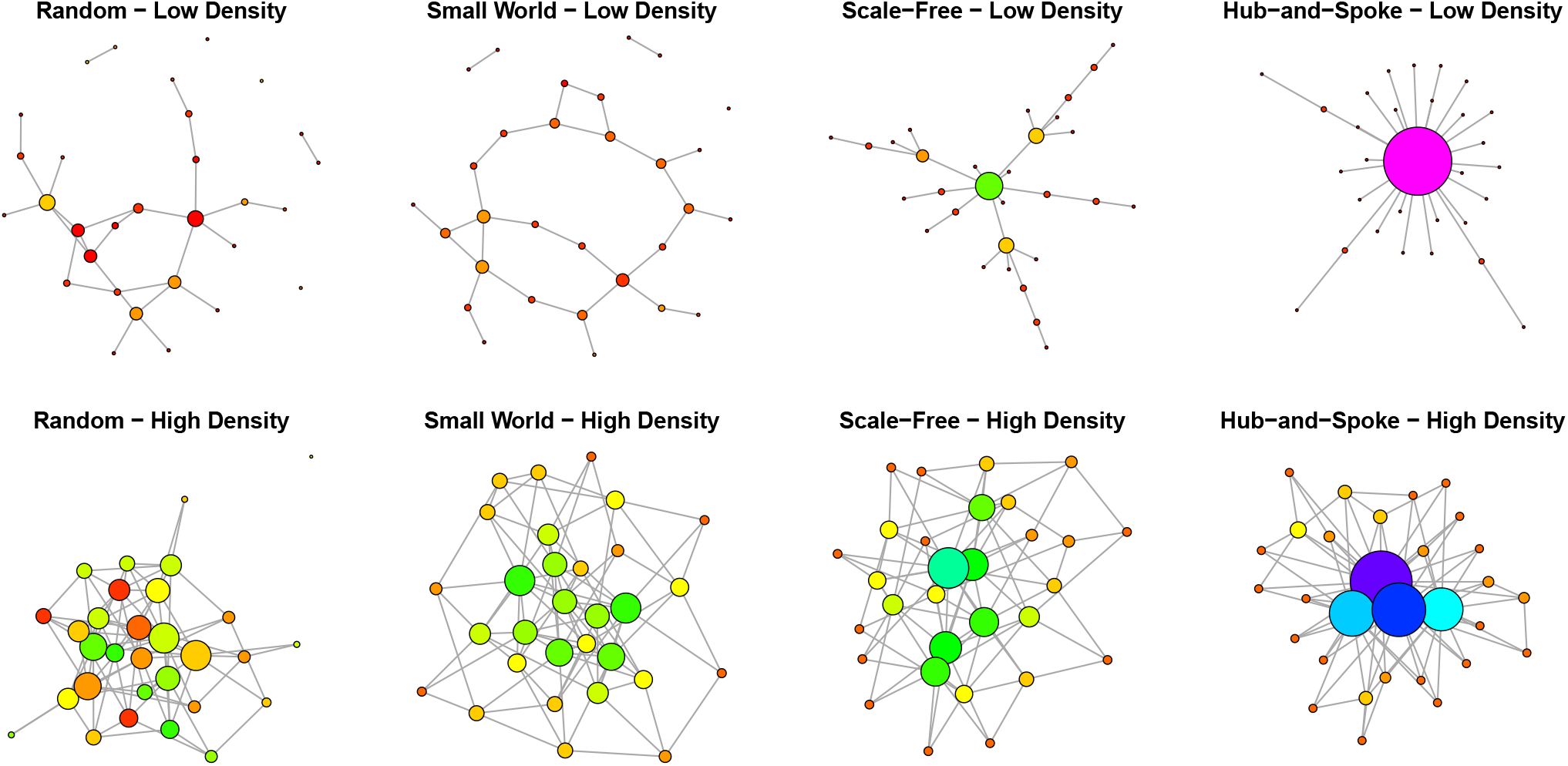
Sample graph topologies simulated using the `igraph` R package. Node size and color indicate degree; larger nodes have higher degree.

### Designing gold-standard networks

The `igraph` package in R was used to simulate gold-standard networks (35). For random networks, the `sample_gnp` function was used (36). For small world networks, the `sample_smallworld` function was used (33). For scale-free and hub-and-spoke networks, the `sample_pa` function was used (Table 1b).

In an effort to create a realistic edge weight distribution from a biological distribution, we used metabolomics data from the CATHeterization GENetics (CATHGEN) biorepository as a starting point (37). The CATHGEN biorepository consists of data from a prospectively-collected clinical study of ~ 10, 000 participants undergoing cardiac catheterization with scheduled annual followup at Duke University Hospital; further details of the study population have previously been published in (37). Measurements of 407 metabolites were available for 136 of these participants, including 68 participants with incident coronary artery disease (CAD) and an equal number of participants without CAD during follow up; further description of this metabolomics study can be found in (38). We used the graphical lasso with the eBIC scoring criterion (hyperparameter *γ* = 0) to estimate a GGM for this dataset. The resulting distribution of the nonzero partial correlations was skewed right with several high outliers (Supplementary Figure S2). We used the histogram of the edge weights as a discrete probability distribution from which to sample edge weights for our simulated networks (bin size 0.01, range −0.32 to 0.76).

To use the weighted adjacency matrix to sample from the multivariate normal distribution, we begin by obtaining a valid precision matrix using Equation 2. For simplicity, we assume *θ_ii_* = *θ_jj_* = 1, which gives the (*i, j*) entry of the precision matrix as:

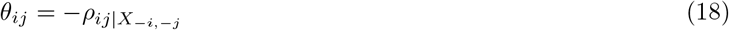

We use this relationship to determine the full precision matrix Θ. Although there is no guarantee that a matrix generated with this approach will be positive definite, we observed positive definite matrices for most of the simulations in this paper. In cases where matrices were not positive definite, we performed a “boosting” step involving adding a small multiple of the magnitude of the minimum eigenvalue to the diagonal of the matrix. For a *p* × *p* precision matrix **A** with minimum eigenvalue 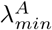, this correction is:

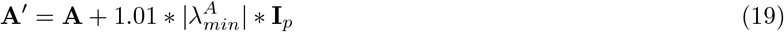

where **I***_p_* is the *p* × *p* identity matrix. This is similar to the approach taken by Tan et. al. in (39).

### Sampling, estimation, dimensionality, and candidate methods

To sample network data from the gold-standard networks, we inverted each estimated precision matrix Θ to find the corresponding covariance matrix ∑, then simulated a sample of size *n* by drawing *X*_1_, …, *X_n_* ~ *MVN*(**0**, ∑). Finally, we estimated precision matrices from this sample in three ways: (i) by applying candidate methods individually (ii) by using a simple mean ensemble model in which each candidate is weighted equally, and (iii) by using the SpiderLearner (Figure 2).

The dimension of a GGM is typically described in terms of the number of samples *n* and the number of predictors *p* included in the model. Importantly, and in contrast to many regression approaches, *p* is not the number of parameters in the model: the precision matrix corresponding to the GGM has *q* = *p* * (*p* – 1)/2 + *p* unique entries that need to be estimated. Because of the quadratic relationship between the number of predictors and the number of parameters to be estimated, dimensionality becomes a major factor in estimation even if a GGM does not include very many predictors. Dimensionalities simulated in this study are shown in Table 1a.

Nine different candidate methods were considered for input to the ensemble algorithm (Figure 2). Candidate Methods 1,2,3,6, and 7 use the `huge` and `huge.select` functions from the R package `huge` with the `glasso` method, which corresponds to the original graphical lasso (7; 3; 6). The difference between these methods is the choice of scoring criterion used in the `huge.select` function to select the tuning parameter (λ in Equation 3). The first criterion is the extended Bayesian information criterion (eBIC), which optimizes a BIC-type quantity tuned by a hyperparameter *γ*, where *γ* = 0 corresponds to a standard BIC measure and *γ* = 0.5 is a typical default value for graphical modeling (12; 40). Candidate Methods 1 and 2 apply this criterion with *γ* = 0 and *γ* = 0.5, respectively. Candidate Method 3 applies a criterion called the rotation information criterion (RIC), which is based on a permutation strategy that generates a null distribution for comparison (12; 7). Candidate Methods 6 and 7 use a criterion called the stability approach to regularization selection (StARS), which is a sub-sampling based approach (12; 41). One of several hyperparameters that can be selected using StARS is `stars.thres`, which relates to the amount of variability that is tolerated across the subsamples (41). Candidate 6 applies the StARS criterion with `stars.thres` = 0.05 and Candidate Method 7 applies it it with `stars.thres` = 0.1 (the default). Candidate Method 4 is the hub graphical lasso, which is an extension of the original graphical lasso that can effectively model hub structures in networks and is implemented in the `hglasso` R package (39). Candidate Method 5 is the MLE, i.e., inverse of the sample covariance as computed with the cov function in base R. Candidate Methods 8 and 9 are similar to Candidate Methods 1 and 2; they also use the original graphical lasso along with an eBIC scoring criterion, but are implemented in the `qgraph` R package (42; 43). A difference between the `qgraph` implementation and the `huge` implementation is in the default range of tuning parameters λ considered. Let λ* be the smallest value of λ that creates an empty graph; `huge` uses a logarithmic sequence of ten candidate λ values between 0.1λ* and λ*, while `qgraph` uses a larger logarithmic sequence of length 100 between 0.01λ* and λ* (7; 43).

We use the following shorthand for these nine methods in the remainder of this paper:

- Candidate Method 1: glasso-ebic-0
- Candidate Method 2: glasso-ebic-0.5
- Candidate Method 3: glasso-ric
- Candidate Method 4: hglasso
- Candidate Method 5: MLE
- Candidate Method 6: glasso-stars-0.05
- Candidate Method 7: glasso-stars-0.1
- Candidate Method 8: qgraph-ebic-0
- Candidate Method 9: qgraph-ebic-0.5

While Candidate Method 5 is typically well-defined in Simulations A-C (barring multicollinearity), it is not in Simulation D, where *n* < *p*. Therefore, Simulation D excludes Candidate Method 5.

### Assessing estimation performance

Once a precision matrix is estimated, we compare it to the original, data-generating, gold-standard precision matrix in order to assess performance. We begin by introducing notation that will be helpful in defining our performance metrics. Let 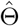 be an estimate of the true *p* × *p* precision matrix Θ, and let 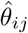 and *θ_ij_* represent the corresponding elements of each. We define the **error matrix** Δ as 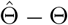, and refer to its *i*, *j^th^* element as:

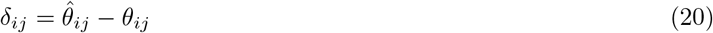

Note that, although the true precision matrix is symmetric, the estimated matrix may not be: a notable example of possible asymmetry is in the graphical lasso algorithm (44). Therefore, we consider every element of the error matrix Δ rather than just upper or lower triangular components when assessing estimation performance.

One area of interest is to assess error in the estimated edge weights in the GGM. Because these edge weights follow directly from the estimated precision matrix, we begin by focusing our efforts on quantifying error in the precision matrix itself. The first metric we use is the based on the size of Δ as assessed by the Frobenius norm:

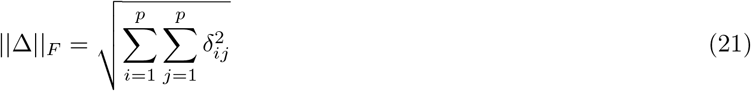

To obtain a quantity that can be compared across topologies, we scale ||Δ||*_F_* by the Frobenius norm of the true precision matrix, ||Θ||*_F_*, defining the **relative Frobenius norm (RFN)** as

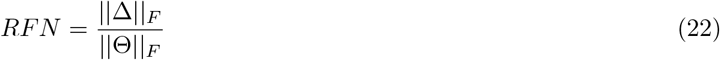

We are interested in the generalizability of the SpiderLearner to independent datasets. For this purpose, we assessed the **out-of-sample log likelihood** of each estimated precision matrix on a new, independent sample of the same size generated from the same gold-standard precision matrix.

Next, we considered bias and MSE for each matrix entry. For *R* simulated replicates of multivariate normal data sampled according to the same gold-standard precision matrix, the **replicate-estimated bias** of element 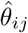 is given by:

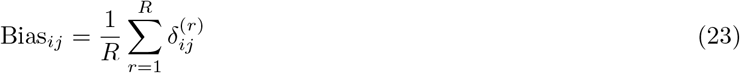

Similarly, the **replicate-estimated mean squared error** of an element *δ_ij_* is given by:

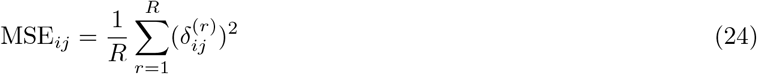

Because most of the candidate models in the ensemble learner are shrinkage methods, we expect that MSE will vary based on the size of each element. Moreover, some methods penalize the diagonal of the precision matrix while others do not. We therefore summarized performance by investigating this element-wise bias and MSE in six categories: (i) the zero elements of the gold-standard matrix, (ii-iv) small, medium, and large entries, corresponding to the bottom quartile, middle 50%, and top quartile of the off-diagonal non-zero matrix elements of the gold-standard matrix, and (v) the diagonal elements of the gold-standard matrix.

In some cases, it may be of interest to assess the performance of a method at estimating the edge set of the GGM, rather than its edge weights. In this case, GGM estimation is essentially a classification problem, where each possible edge (*i, j*) is classified as either included in the network or excluded from the network. For this purpose, it is necessary to classify edges in some way. In Simulations A-C, where the MLE is included as a candidate learner, it is likely that the estimated GGM will be completely connected as the MLE will not contain entries that are exactly zero. For these three simulations, we therefore construct a sparser graph by thresholding based on the significance of the Fisher-transformed partial correlation coefficient (see Supplement for details). We control the FWER at level *α* = 0.05 with a Bonferroni correction (45).

In Simulation D, the Fisher-transformed partial correlation is not well-defined (see Supplement). We therefore classify any non-zero partial correlation coefficient as an edge. Because the MLE is not included in Simulation D, this classification yields a graph that is not completely connected. This methodology is described in detail in the Supplement.

As an additional measure, we calculated the **matrix RV coefficient**, an analogue of a correlation coefficient, between estimated and gold-standard matrices, as implemented in the R package `MatrixCorrelation` (46; 47).

## Results

We conducted 100 iterations for each of the eight network topologies in each of Simulations A-D. Results for Simulation A and Simulation D are presented here; results for Simulation B and Simulation C are presented in the Supplement. To provide some perspective of computation times, a table of runtimes for various values of *n* and *p* is available in Supplementary Table S1.

### Simulation A: Sample size >> number of features, parameters estimated (*n* >> *p,q*)

Ensemble weights for Simulation A are shown in Table 2a. The SpiderLearner algorithm selected at least three different methods to have nonzero weights for each topology, demonstrating that combining multiple candidate algorithms is indeed important from a likelihood-based loss perspective. For every topology, `qgraph-ebic-0` and the inverse sample covariance (i.e., MLE) were included in the combination, although the weights varied broadly by topology, with `qgraph-ebic-0` weights ranging from 0.03 for the low-density hub-and-spoke topology to 0.42 for the low-density scale-free topology and the MLE weights ranging from 0.28 for the low-density random graph topology to 0.59 for the high-density random graph topology. The `hub glasso` was included in seven out of eight topologies (excluding the low-density scale-free topology), again with broadly varying weights (0.10-0.63). The `glasso-ebic-0` was selected for minor contributions in the low-density scale-free case(0.33) and the high-density scale-free case (0.08). The `glasso-ric`,`glasso-stars-0.05` and `glasso-stars-0.1` methods were weighted zero for all topologies.

**Table 2:**
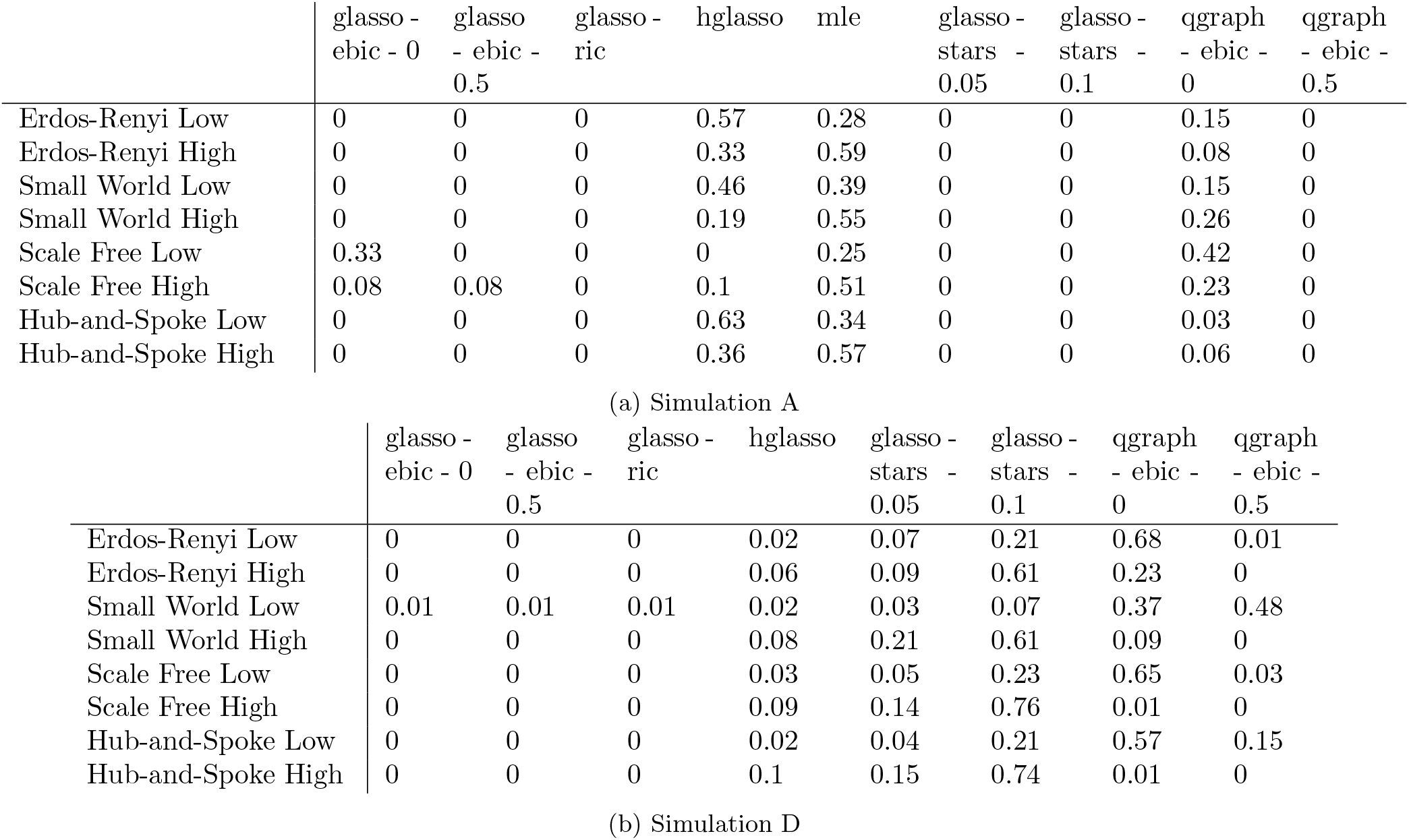
Average weight for each method as selected by SpiderLearner in N=100 simulations.

Results for the relative Frobenius norm of the error matrix are shown in Figure 4a. It can be easily seen that the performance of each method varied highly according to this metric, emphasizing the importance of our approach. The SpiderLearner performed better than the individual candidates and better than the simple mean ensemble model across all settings considered. The performance as assessed by out-of-sample log likelihood can be seen in Figure 4b. Again, the SpiderLearner performed well; we note that variability of the out-of-sample log likelihood was not as high as that of the RFN.

**Figure 4:**
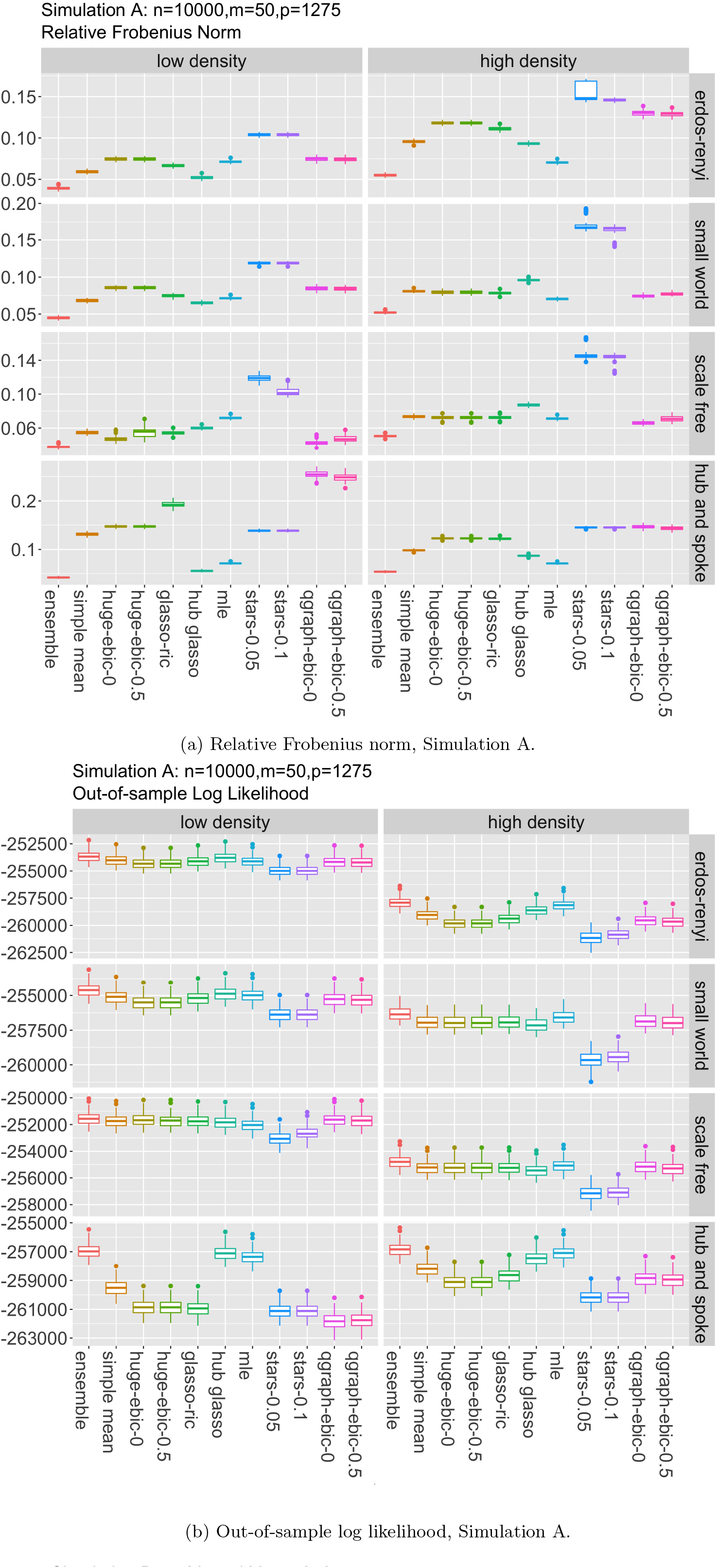
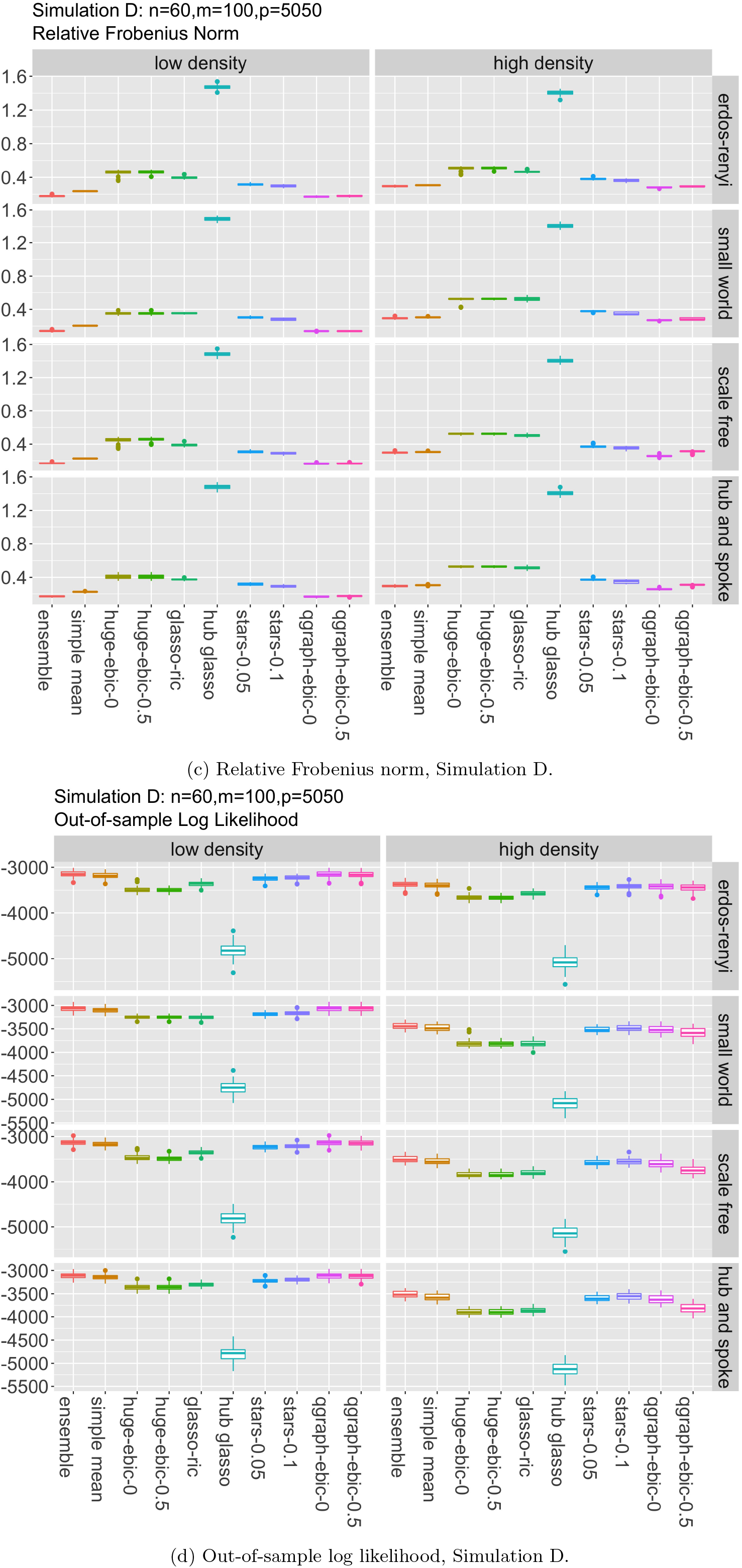
Simulation A and D results show that the SpiderLearner ensemble model is able to outperform or match the performance of every other candidate included in the model, and exceeds the performance of a simple mean of the candidates. Relative Frobenius norm results demonstrate the ability of the algorithm to accurately estimate the precision matrix entries, while out-of-sample log likelihood shows that the estimated precision matrix is not overfit by the SpiderLearner.

The element-wise bias and MSE for the five entry categories (zero, small, medium, large, and diagonal) are shown in Supplementary Figure S4. For the SpiderLearner as well as all the candidate methods, bias varied by entry category. The MLE showed the smallest bias in each case, which is logical given that it was the only non-shrinkage method employed. The SpiderLearner performed better than or comparably to the remainder of the algorithms in terms of the magnitude of bias, while having the added benefit of smaller variability of bias for most elements. The exception was for the true zero elements, in which the SpiderLearner incorporated the MLE and had a higher variability accordingly. A similar pattern was observed for MSE.

Sensitivity and specificity of each method in Simulation A are shown in Supplementary Table S2 and S3, respectively. The SpiderLearner was more sensitive than some candidate methods and less sensitive than others; it was more sensitive than the simple mean in every case. It had perfect specificity (as did most candidate methods), selecting no false positives. This observation may be due to the conservative nature of our threshold, which is based on a Bonferroni correction (see Supplement).

### Simulation D: Sample size < number of features << number of parameters estimated (*n* < *p* << *q*)

Ensemble weights for Simulation D are shown in Table 2b. The SpiderLearner algorithm selected at least four of the candidate methods in every case. Interestingly, the `glasso-ebic-0`, `glasso-ebic-0.5`, and `glasso-ric` contributed the least to the ensemble in Simulation D after being important players in Simulation A. On the other hand, `glasso-stars-0.05` and `glasso-stars-0.1` both contributed to the ensemble in all eight cases in Simulation D, but they were not selected in any case in Simulation A. The hub graphical lasso was a highly-weighted candidate in most cases in Simulation A, but had very low weights in Simulation D. These observations are further evidence of the important of considering multiple methods when estimating a GGM: the performance (in the log-likelihood sense) of estimates from different methods varies broadly based on the characteristics of the true underlying network.

Results for the RFN in Simulation D are shown in Figure 4c. We generally saw the SpiderLearner performing comparably to the `qgraph-ebic-0` candidate method and `qgraph-ebic-0.5`, two methods which were typically highly weighted in the ensemble (Table 2b). The out-of-sample log likelihood performance is shown in Figure 4d. The SpiderLearner again performed well when compared to the remainder of the methods. The hub graphical lasso had notably lower out-of-sample log likelihood than the other candidates, suggesting overfitting in this setting.

Bias and RMSE for Simulation D are shown in Supplementary Figure S10. In Simulation D, the hub graphical lasso clearly outperformed all the other methods in terms of bias, while suffering a large MSE (i.e., high variance). The SpiderLearner was able to detect this tradeoff and avoid excessive variance by assigning a low weight to the hub graphical lasso. Aside from the hub graphical lasso, bias and MSE were comparable across the SpiderLearner, the simple mean, and the remaining candidate methods for the zero, small, medium, and large entries. Bias differed for the diagonal entries as some of the candidate methods applied a shrinkage penalty to the diagonal, while some did not.

In Simulation D, candidate methods either had a moderate sensitivity or a very low sensitivity; the SpiderLearner fell into the moderate category, with sensitivity around 0.6. (Supplementary Table S10). Many methods with low sensitivity selected empty graphs, possibly due to the small sample size relative to the number of predictors in this simulation setting. Specificity was similarly bimodal, with the SpiderLearner, the simple mean, and the hub graphical lasso having a specificity around 0.45, while other methods had a specificity of near 1 (Supplementary Table S11).

### The MLE as the estimator for the precision matrix

In simulation settings A,B, and C, the sample size *n* is larger than the number of predictors *p* in the model, meaning that the sample covariance matrix is non-singular, except in the case of multicollinearity. The sample covariance matrix is the MLE for the population covariance matrix, and because inversion of a non-singular matrix is a continuous function, the inverted sample covariance matrix is the MLE for the population precision matrix (48; 49). Notably, that the likelihood-based SpiderLearner model selects models other than the MLE, and that other individual regularized algorithms perform better than the MLE according to the relative Frobenius norm, matrix RV coefficient, and out-of-sample likelihood. We hypothesized that this phenomenon was related to the sparsity of the underlying network. To investigate, we ran the SpiderLearner algorithm on an Erdös-Renyí random graph with a variety of densities (0.05, 0.1, 0.25, 0.5, 0.75, 1) with 30 iterations for each density. As hypothesized, the weight of the MLE in the ensemble model increases with the density of the graph, as shown in Figure 5. These results suggest that even though the ensemble loss function does not incorporate a shrinkage penalty, it is still advantageous from the likelihood-based perspective to shrink estimates of small precision matrix entries to zero in the case where the population precision matrix is sparse. The takeaway is that shrinkage methods can improve out-of-sample performance even in low-dimensional cases, which is consistent with results observed in the original LASSO publication (50).

**Figure 5:**
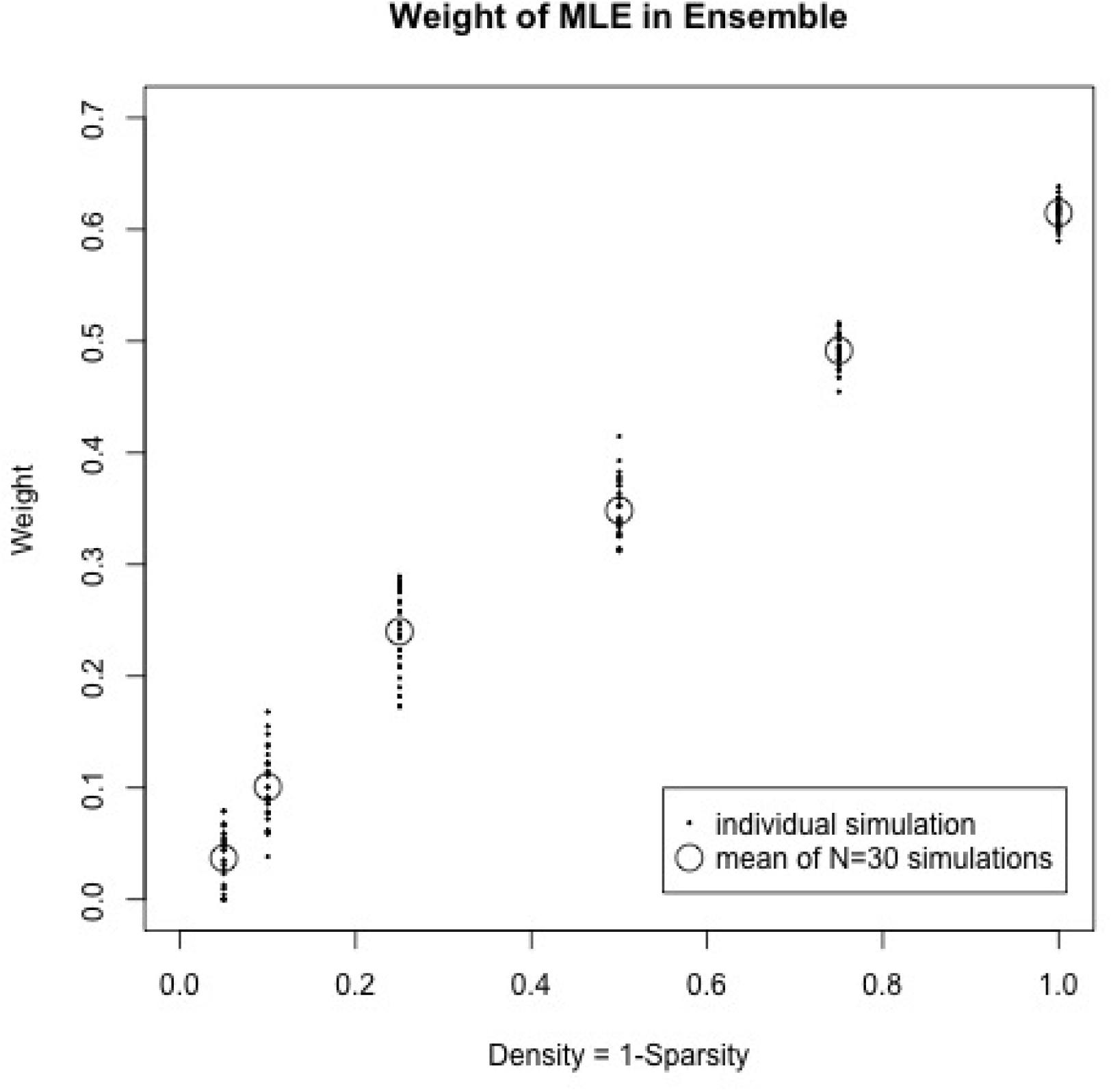
Using a random graph topology with six different densities (0.05, 0.1, 0.25, 0.5, 0.75, and 1), we explored the relationship of the weight of the MLE in the ensemble model with the graph density. In sparse graphs, the MLE is not weighted heavily by the algorithm; as density increases, the MLE begins to dominate the contribution to the convex combination.

### Choice of *K*

A practical question in this methodology is how to select *K* in the *K*-fold cross-validation. Higher values of *K* give more training data, meaning the estimates of the candidate precision matrices Θ_1_, …, Θ*_M_* are more accurate and more precise; however, less data are available to estimate the out-of-sample log likelihood on the left-out test data, meaning estimates of ***α*** will suffer higher bias and variance. Lower values of *K* give less training data and more out-of-sample data, but it is not immediately clear that this causes the reverse problem: quality of estimates of ***α*** depend both on the amount of out-of-sample data as well as the quality of the estimates of Θ_1_, …, Θ*_M_*.

It is apparent that there is a complex “Θ-*α* tradeoff” underlying our method, making it challenging to recommend a particular choice of *K* without further investigation. For these reasons, we conducted a simple simulation study to assess the impact of the use of different values of *K*. We used the high-density Erdös-Renyí random graph topology as a gold standard network, generated 100 samples of size *n* = 150 on *p* = 50 predictors (*q* = 1275 parameters to be estimated), and ran the SpiderLearner algorithm for *K* ∈ {2, 5, 10, 15, 20, 30}. We then calculated (i) the element-wise standard deviation of the estimated precision matrix 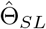, (ii) the element-wise bias of 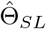, and (iii) the variability of the selected coefficients 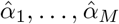 for each value of *K*. Because the library includes shrinkage methods, we calculated summary measures for (i) and (ii) in four categories of the true matrix: zeros, bottom 10 percent of non-zero entries, middle 80 percent of non-zero entries, and top 10 percent of non-zero entries. Because the true matrix is symmetric, we only assessed diagonal and lower triangular elements.

Supplementary Figure S11 shows that the element-wise standard deviation of 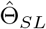 increases slightly with increases in *K*, but that most of the change happens when moving from *K* = 2 to *K* = 5 and from *K* = 5 to *K* = 10. Supplementary Figure S12 shows that entry-wise bias decreases substantially as *K* increases for medium and large entries; although it slightly increases as *K* increases for zero entries and for small entries, the magnitude of these increases is small compared to the decrease in bias for the medium and large entries. Supplementary Figure S13 shows that the variability of the weights 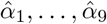 is similar across all considered values of *K* ≥ 5. These results lead us to suggest that *K* = 10 is a good choice, with *K* = 5 as an option for large datasets if computing time is a limitation.

### Selection of candidate GGM library

Existing Super Learner literature suggests that a broad and varied library of candidate learners is beneficial, and that overfitting or highly variable estimates are typically not observed consequences of a large library size, although cross-validating the Super Learner itself is recommended as a best practice (15; 24; 51). Practical limitations to library size include computation time and interpretability. We investigated the sensitivity of model results to the library size and content in 100 simulated datasets of size *n* = 1000, *p* = 50 using the high-density scale-free graph topology. We used three different libraries: (i) a small baseline library consisting of the hub graphical lasso and the MLE, chosen due to the ability of the hub graphical lasso to model the scale-free topology and the generally favorable properties of the MLE, (ii) a medium library consisting of the small library along with the `huge-ebic-glasso` method with *γ* = 0 and *γ* = 0.5, and (iii) a large library consisting of the nine methods used in Simulations A-C. Our results indicate that while the large library provides the best fit, the results from the medium and small library do not differ substantially as a whole in this case (Supplementary Figure S14). In addition to this simulation setting, we further explored the sensitivity of the estimated network to the library selection in a real data example, in which we observed that some libraries yield similar results while others differ (Supplementary Table S12, Supplementary Figure S16).

### Application: Ovarian cancer risk modeling

#### Datasets

To demonstrate the application of our method to real data, we used sixteen ovarian cancer gene expression datasets from the Curated Ovarian Cancer collection of Ganzfried et. al. (20). One of the datasets was used to train a SpiderLearner model and develop a network-based risk score from the resulting network; the other fifteen were used as independent validation datasets to evaluate the risk score performance (Figure 6). The training dataset consists of 260 late-stage ovarian cancer patients with gene expression data for 20106 genes, obtained via microarray experiments by Yoshihara et al. (19) (“Yoshihara dataset”). Characteristics of the Yoshihara dataset have been previously described in (19), where it is referred to as Japanese data set A. Briefly, the study conducted by Yoshihara et al. included participants with advanced stage high-grade serous ovarian cancer who underwent debulking surgery followed by chemotherapy, with followup for up to ten years. Yoshihara et al. assessed overall survival was assessed as the time from the primary surgery to death due to ovarian cancer. 131 of the 260 patients were living at the end of the study; we treated these patients’ outcomes as right-censored. Basic characteristics of all sixteen datasets are shown in Table 3. For details regarding the 15 validation datasets, we refer the reader to the original publications, also shown in Table 3. In the application below, we used the nine candidate GGM estimation methods described in Figure 2 as the SpiderLearner library, with *K* = 10-fold cross-validation. All data are publicly available through the R package `curatedOvarianData` (via Bioconductor), and code to reproduce the application workflow is available at https://github.com/katehoffshutta/SpiderLearnerWorkflow.

**Figure 6:**
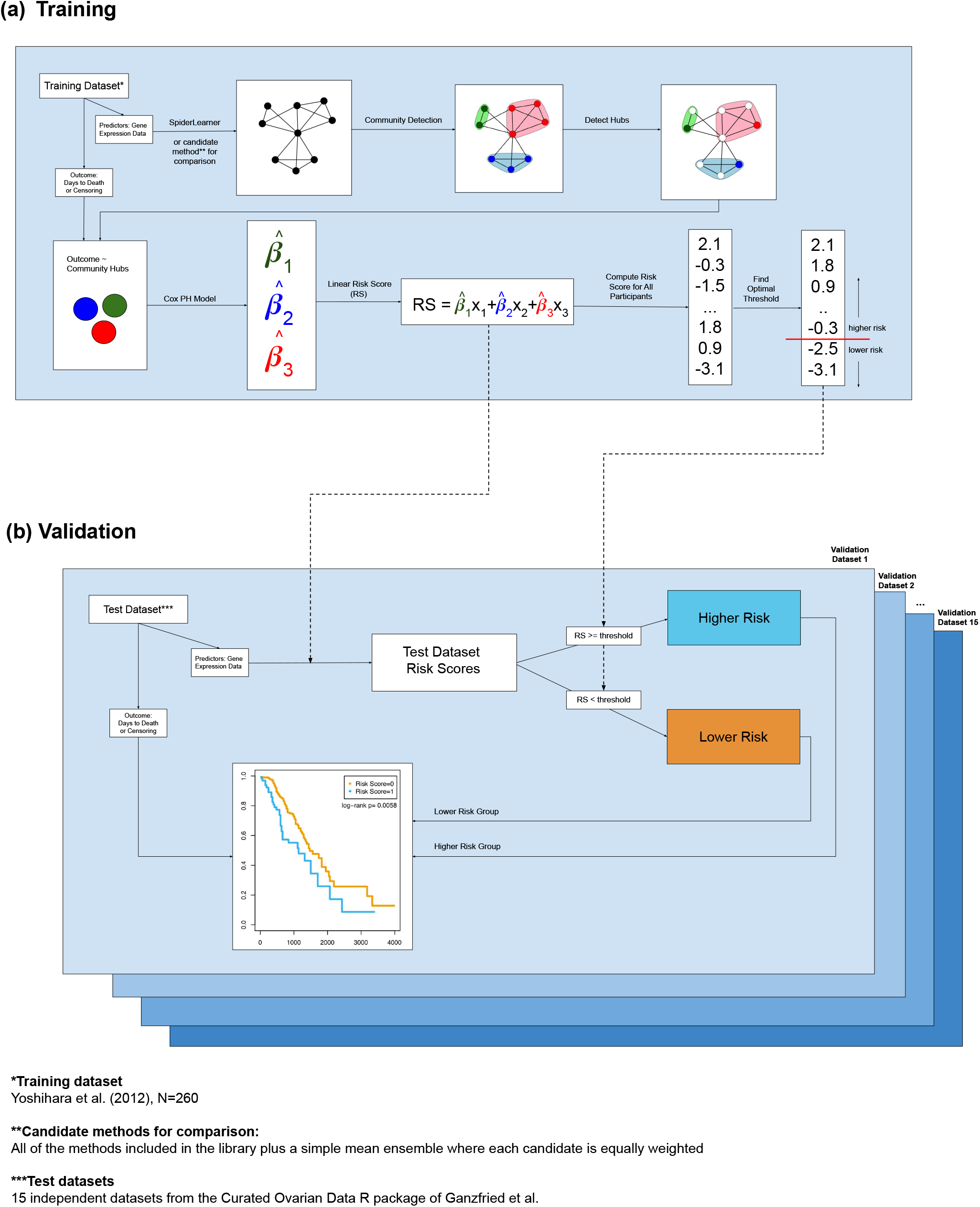
Workflow for training and validating the network-based risk score.

**Table 3:** Basic characteristics and references for the 16 ovarian cancer datasets used in the SpiderLearner application.

| Dataset | Platform ID | N | Age: Mean(SD)<br>*missing* | Tumor<br>Stage<br>(% < 4)<br>*missing* | Summary<br>Stage<br>(% Late)<br>*missing* | Summary<br>Grade<br>High (%)<br>*missing* | Validated<br>in<br>KM<br>Models | Validated<br>in CoxPH<br>Models |
| --- | --- | --- | --- | --- | --- | --- | --- | --- |
| GSE32062.GPL6480 (19) | hgug4112a | 260 | - | 78 | 100 | 50 | - | - |
| E.MTAB.386 (52) | illuminaHumanv2 | 129 | 60.71(14.24) | 85 | 99 | 1 | no | yes |
| GSE13876 (53) | OperonHumanV3 | 157 | 57.95(12.39) | - | 100 | 54 13 | yes | yes |
| GSE14764 (54) | hgu133a | 80 | - | 98 | 89 | 68 | yes | yes |
| GSE17260 (55) | hgug4112a | 110 | - | 80 | 100 | 39 | no | no |
| GSE18520 (56) | hgu133plus2 | 63 | - | 100 10 | 84 10 | 84 10 | no | no |
| GSE19829.GPL570 (57) | hgu133plus2 | 28 | - | - | - | - | yes | no |
| GSE19829.GPL8300 (57) | hgu95av2 | 42 | - | - | - | - | no | no |
| GSE26712 (58) | hgu133a | 195 | 61.54(11.86) 13 | 80 13 | 95 10 | 95 10 | no | no |
| GSE30009 (59) | NA | 103 | 62.45(11.14) | 80 | 100 | 89 2 | no | no |
| GSE30161 (60) | hgu133plus2 | 58 | 62.57(10.61) | 91 | 100 | 57 4 | no | yes |
| GSE32063 (19) | hgug4112a | 40 | - | 78 | 100 | 42 | no | no |
| GSE9891 (61) | hgu133plus2 | 285 | 59.62(10.59) 3 | 92 3 | 84 3 | 57 6 | yes | yes |
| PMID17290060 (62) | hgu133a | 117 | - | 85 1 | 98 1 | 49 3 | yes | no |
| PMID19318476 (63) | hgu133a | 42 | 61.46(10.61) 1 | 76 1 | 93 1 | 57 1 | yes | no |
| TCGA (64) | hthgu133a | 578 | 59.7(11.56) 10 | 85 15 | 90 15 | 83 23 | no | no |

#### Workflow and results

In (19), Yoshihara et al. present a 126-gene signature of high-risk ovarian cancer based on overall survival, defined as time from primary surgery to death or loss-to-followup. To investigate the relationships between the genes in this signature, we used SpiderLearner to estimate a GGM for this study setting. We extracted 116 of the genes presented in (19) from an example dataset in the `curatedOvarianData` R package. The weights selected by SpiderLearner were 0.69 for `hglasso`, 0.13 for `huge-ebic-0`, 0.12 for `qgraph-ebic-0`, 0.05 for the MLE, and zero for the remainder of the candidate algorithms.

Community detection is a useful way to identify clusters in graphical models. We applied the `cluster_walktrap` community detection algorithm as implemented in the `igraph` R package to detect communities in the SpiderLearner-estimated GGM as well as the GGMs estimated by the nine candidate algorithms and the simple mean (65; 35). The `cluster_walktrap` algorithm requires the choice of a step size for the random walk. For each network, we selected the step size between 1 and 10 that maximized the overall modularity of the network. Estimates for the nine candidate methods are shown in Figure 7a. The community structure varies notably across methods, further motivating the use of our ensemble method.

**Figure 7:**
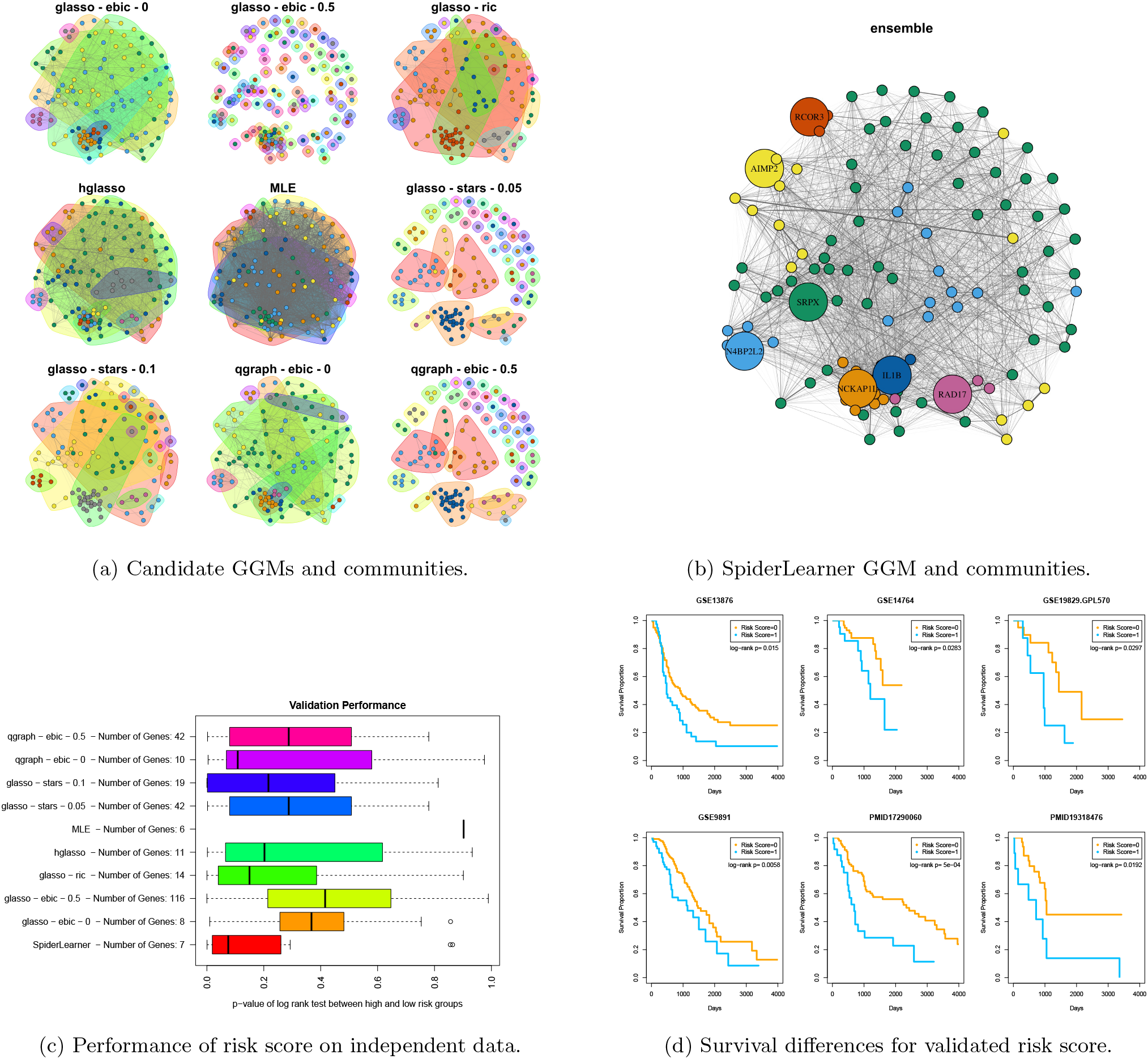
(a) Estimated GGMs and communities for 116 of the genes present in the Yoshihara 126-gene signature for each of the candidate methods in the SpiderLearner library. Vertices of the same color belong to the same community. Estimated community structure varied widely by candidate method. (b) SpiderLearner-estimated network, including the seven local hubs used in the risk score. (c) Boxplots of the validation log rank test *p*-value testing the null hypothesis of no difference between high-risk and low-risk estimates as defined by the hub-based risk score versus the two-sided alternative. The ensemble model shows a better validation performance and uses a more parsimonious risk score model than other candidates, requiring only seven genes as predictors. (d) The SpiderLearner-based risk score includes the seven labeled genes in (b) as predictors. For the six of 15 validation datasets in which the SpiderLearner-based risk score was successfully validated, Kaplan-Meier estimates for the low-risk (risk score = 0) and high-risk (risk-score=1) groups are shown in (d) along with the *p*-value of the log-rank test comparing the two curves.

To derive biological insight from the detected communities, we developed a network-based risk score utilizing topological characteristics of the estimated GGM to identify a set of genes with which to predict overall survival. We closely followed the approach of (19) in developing their ovarian cancer prognostic index. (19) began by using a penalized Cox proportional hazards (Cox PH) model to obtain regression coefficients for each of the 126 genes. Next, the authors calculated a prognostic index as follows:

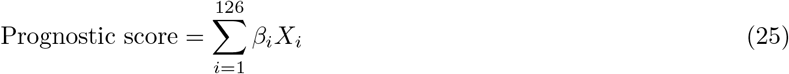

where *β_i_* was the regression coefficient for gene *i* in the penalized Cox PH model and *X_i_* was its centered and standardized gene expression value. Finally, (19) determined the optimal threshold value of their prognostic index by (i) assigning patients to a high-risk or low-risk group based on a proposed threshold, (ii) calculating the *p*-value of a log-rank test for difference in overall survival between the high-risk and low-risk group, and (iii) repeating this process for a number of thresholds and finding a threshold that minimized the *p*-value in (ii).

The workflow that we applied is in the same spirit, and is shown in Figure 6a. Rather than using all 126 genes to produce the score, we aimed to find more a more parsimonious score by leveraging the network structure of the Yoshihara dataset to select a subset of relevant genes. For each candidate approach, the simple mean, and the SpiderLearner, we began by identifying the gene in each community with the highest hub score by applying the `hub_score` function of the `igraph` R package to the adjacency matrix of the estimated GGM (35; 66). Hub scores reflect how influential nodes are based on the eigendecomposition of the weighted adjacency matrix of a graph, with a higher hub score corresponding to more influence (66). Earlier work in bipartite networks of SNPs and genes has demonstrated that hubs within communities are enriched for disease-associated SNPs (67). We hypothesized that local hubs in GGM communities might have similar functional leverage and therefore be useful predictors of ovarian cancer outcomes.

To develop the risk score, we fit Cox PH models regressing days to death on the local hubs in each network using the `survival` package in R (68). For GGM estimation method *m* with local hubs *x*_1_, …, *x_p_*, we denote the corresponding Cox PH model coefficients as 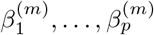. Following the development of the prognostic index in (19), the risk score for patient *i* according to method *m* was calculated as

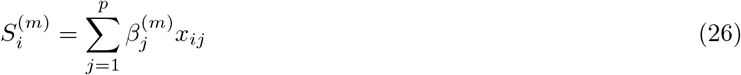

where *x_ij_* is the centered, standardized expression level of gene *j* for person *i*.

We next mapped the score in Equation 26 to a binary indicator of high risk or low risk by establishing a threshold point. As in (19), we selected an optimal threshold for this score by testing a grid of threshold values and selecting the value attaining largest separation between the estimated Kaplan-Meier survival curves of the high-risk and low-risk groups in the Yoshihara dataset, as measured by the lowest *p*-value according to a log-rank test of the difference. For the ensemble method, this threshold was 0.461 (log-rank test *p* = 3.7 * 10^−10^).

The framework for validation of the risk score is shown in Figure 6b. Each of the ten total risk scores and associated thresholds for distinguishing low- vs. high-risk participants was evaluated in 15 independent ovarian cancer datasets comprised of data available from (20) that contained information on the outcome (days to death) as well as vital status. A risk score was considered to validate if the log-rank *p*-value between the Kaplan-Meier estimates for the high-risk and low-risk groups was less than 0.05. Figure 7c shows the distribution of the validation *p*-values across the 15 datasets for the risk score developed from the ensemble network as well as from the nine candidate networks. In some cases, the optimal threshold score determined from the training dataset was such that there were insufficient samples above and below the threshold to perform a log-rank test in the validation dataset; these cases are omitted from the boxplot (MLE: N=14 of 15 studies, ensemble: N=1, `hglasso`: N=1, `glasso-ebic-0`: N=1). Table 4a shows further detail about the validation. Notably, in the case of `glasso-ebic-0.5`, an empty network was selected. Consequently, every gene formed its own community and all 116 genes were required to construct the risk score. We can thus use the `glasso-ebic-0.5` case to benchmark the ensemble method vs. a naive approach in which the network structure is not leveraged to construct the risk score. Figure 7c shows that the ensemble approach provides a considerable gain.

**Table 4:**
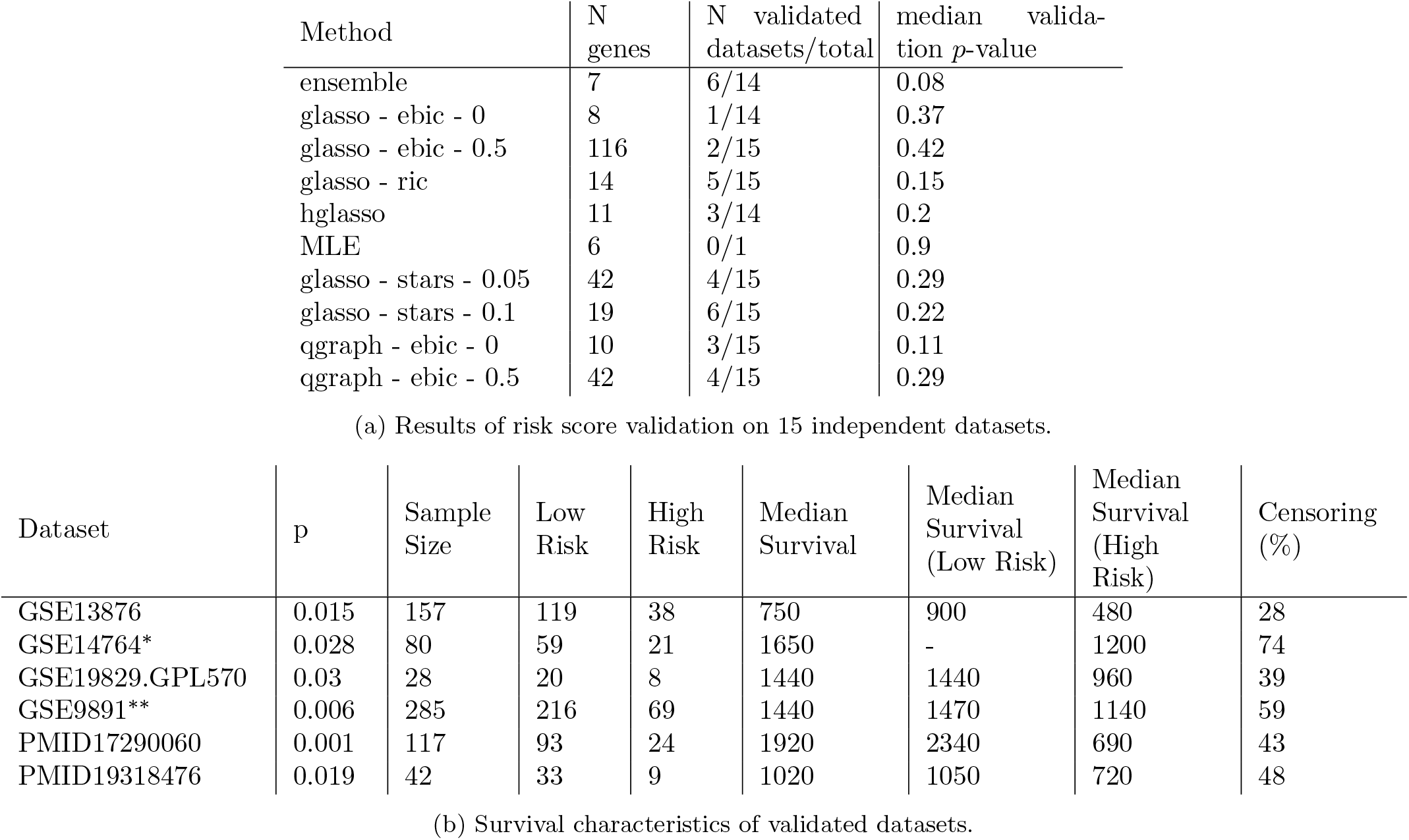
Results from assessing risk score in validation datasets demonstrate that the ensemble network produced the most robust risk score from the training dataset. Median survival (in days to death) is estimated from the Kaplan-Meier curves for the six datasets in which the seven-gene risk score validated. *In the GSE14764 study, more than half of the low-risk participants survived past the end of the study, so the median survival was undefined in this group. The time of last observation in the low-risk group was 1590 days. ** Vital status missing for 3 participants in GSE9891

We note that the SpiderLearner risk score model has two important advantages over the risk score developed by each candidate method. First, it is a much more parsimonious model, requiring only seven predictors to develop a risk score that validated in 6 of the 15 validation datasets, whereas the only other method achieving this performance required 42 genes to do so. Second, it has the lowest median validation *p*-value across the 15 validation datasets, and that median approaches nominal significance (SpiderLearner median validation *p* = 0.08). This result indicates better risk prediction even among those datasets in which the risk score did not validate according to the *p* < 0.05 criterion. In an effort to steer clear of overvaluing *p*-value thresholds, we emphasize that these overall, non-thresholded, results also reinforce the robustness of the SpiderLearner risk-score. Kaplan-Meier plots for the SpiderLearner-based risk score on the six datasets in which it validated are shown in Figure 7d. Similar plots for all 15 datasets are available in Supplementary Figure S17. Median survival times in the low-risk and high-risk group differ substantially (Table 4b), suggesting our method is capable of producing findings with clinical relevance.

In order to explore the influence of available clinical covariates on the relationship between our risk score and survival, we performed an additional analysis involving Cox PH models. We first estimated the unadjusted hazard ratio of the risk score with a model including only the risk score as a covariate. Next, we estimated the adjusted hazard ratio, adjusting for the following covariates where available: age at time of pathological diagnosis, summary grade (low, high), summary stage (early, late), and tumor stage (< 4, 4). The unadjusted hazard ratio for the risk score was significant in five of the 15 validation sets; in these five cases, the adjusted hazard ratio was also significant (Table 5, Supplementary Figure S18). These results indicate that our risk score provides additional prognostic information above and beyond that contained in these clinical characteristics. Further, the median *p*-value of the hazard ratio of the SpiderLearner risk score was comparable to that of the best candidate methods, even though it used fewer predictors to generate the score (Supplementary Figure S19).

**Table 5:**
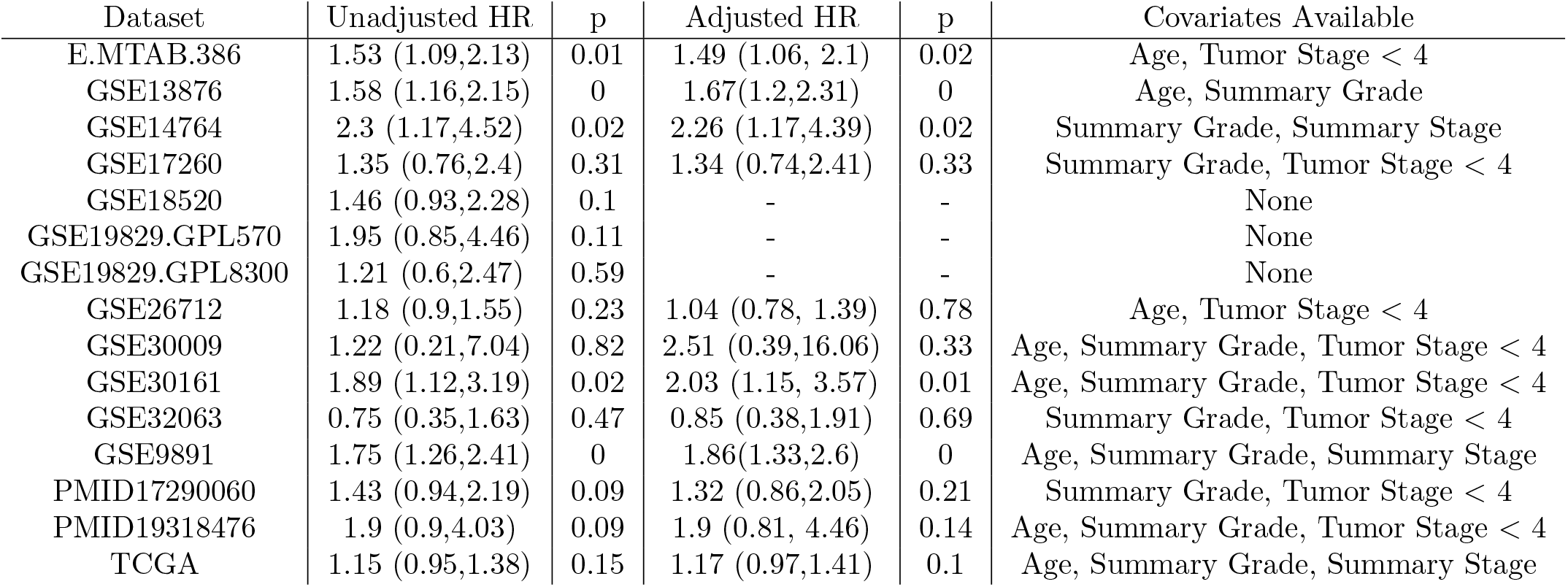
Estimated unadjusted and adjusted hazard ratios and 95% confidence intervals for the SpiderLearner risk score. Confidence intervals represent the exponentiated the endpoints of a Wald-type 95% confidence interval for the log hazard ratio.

Inspection of Table 3 shows no substantial differences in age, tumor stage, summary grade, or summary stage between the datasets in which the risk score validated and those in which it did not. We note that 5 of the 6 validated datasets in which the risk score validated used the hgu133a (Affymetrix Human Genome U133) platform or the hgu133plus2 (Affymetrix Human Genome U133 Plus 2.0) platform, while 8 of the 15 used one or the other of these. The association between platform and validated status was not statistically significant (Fisher’s exact test, p = 0.12).

## Discussion

In this work, we establish SpiderLearner, an ensemble method for estimating a Gaussian graphical model (GGM) from a convex combination of precision matrices estimated using a broad range of existing open-source candidate methods. In a wide variety of simulation settings, SpiderLearner consistently performed comparably to or better than each of the candidate methods according to a variety of metrics, including relative Frobenius norm of the error matrix, matrix RV coefficient, and element-wise MSE. Importantly, some of the individual candidate methods performed quite poorly; since a researcher’s best option *a priori* is to simply choose one of the candidate methods at will, our ensemble method provides a considerable advantage for practical use.

New methods for GGM estimation are being continually developed and assessed. For example, Lartigue et al (2020) conduct an extensive simulation study on GGM estimation for small sample sizes and present a composite procedure that uses a likelihood criterion to select a GGM (69). Methods such as these that are specific to the particular research settings such as the small-sample case are areas for further development. An advantage of SpiderLearner is that such methods, when developed, can be included as candidate models in the ensemble library.

We demonstrated the practical utility of our approach by modeling the network-level interactions of genes belonging to a previously published 126-gene signature of high-risk ovarian cancer (19). The seven genes selected by the ensemble model for inclusion in the risk score are *AIMP2, NCKAP1L, SRPX, NĄBP2L2,IL1B,RAD17*, and *RCOR3* (Figure 7b). All seven of these genes have important biological function, with experimental evidence linking their expression levels to processes such as cell proliferation and immune system function that have implications in the study of the development, progression, and treatment of cancer.

*AIMP2* is an important tumor suppressor gene, and its splice variant *AIMP2-DX2* has been shown to be an effective pharmaceutical target in chemotherapy-resistant ovarian cancer (70).

Recent work demonstrated that *in vitro* overexpression of *SRPX* resulted in increased ovarian cancer cell invasion activity, while shRNA reduction of *SRPX* mRNA led to a decrease (71).

*RAD17* encodes a protein that is related to checkpoint signalling in the cell cycle; RAD17 expression is oscillatory, and engineered stabilization of RAD17 resulted in disrupted checkpoint signalling and consequent diminished re-entry into the cell cycle (72).

*RCOR3* encodes a protein called CoREST/REST corepressor 3 and is a paralog of RCOR1, a protein which works together with lysine-specific demethylase 1 (LSD1) in epigenetic regulation of cell fates (73). Upadhyay et al. (2014) demonstrate that RCOR3 is recruited to target genes by LSD1 along with a protein called growth factor independent 1B transcriptional repressor (GFI1B), decreasing histone demethylation and thus de-repressing target gene expression. LSD1 is known to repress tumor suppressor gene expression in oncogenesis, and it is suggested that an increase in RCOR3 expression could attenuate this contribution to oncogenesis.

*N4BP2L2* encodes a protein known as (N4BP2L2 full name) or as phosphonoformate immunoassociated protein 5 (PFAAP5) (74). There is evidence that N4BP2L2 is involved in neutrophil deficiency (neutropenia), participating in transcriptional regulation of a neutrophil production pathway (74).

*IL1B* encodes the interleukin-1*β* protein, which has been shown to be higher in serum and plasma of ovarian cancer patients relative to healthy women and has been implicated in important signaling cascades, including the p38/JNK pathway and the NF-*κ*B pathway (75). *NCKAP1L* has recently been identified as a novel tumor micro environment-related biomarker in luminal breast cancer, but has not been previously studied extensively in relation to ovarian cancer (76). *IL1B* and *NCKAP1L* are both members of a number of interesting GO biological processes, including the regulation of phagocytosis, vascular EGFR regulation, neutrophil chemotaxis and migration, granulocyte chemotaxis, and regulation of T-cell, interleukin-6, lymphocyte, and mononuclear cell proliferation.

Recent advances have been proposed to improve the applicability and reproducibility of network estimation methods. (77) propose a Monte Carlo-based method for generating confidence intervals for network statistics, allowing a researcher to assess whether a network property such as edge presence or node centrality differs from that expected by random chance. (78) present a bootstrap-based approach which allows researchers to investigate the variability of an estimated network. (79) develop a network meta-analysis framework that permits integration of estimated networks across multiple studies. Each of these methods can be in theory be applied to GGMs, but rely on the use of an initial estimation algorithm. Consequently, results will still remain sensitive to the many choices that the researcher must make during the estimation process. Our method can thus complement the advances described above, potentially contributing to improved reproducibility and generalizability in GGM estimation.

A limitation of this approach is its time-consuming nature; for *K*-fold cross validation with *M* candidate models, the time cost of estimating the ensemble model would be about *M*(*K* + 1) times the cost of estimating just one candidate model (assuming all candidates take roughly the same amount of time). Moreover, the number of model parameters to be estimated by each candidate model grows quadratically with the number of predictors included in the network, meaning that the computational cost of the ensemble model can quickly become substantial for larger predictor sets. Because model fitting in each fold is independent, parallelization is a good solution to this problem when multiple cores are available. We have implemented parallel processing in the SpiderLearner code to help reduce runtime.

A second limitation lies in the rigidity of the convex combination of precision matrices. The same coefficient is applied to every element of each precision matrix in the current ensemble model formulation. A more flexible extension could address this limitation by partitioning matrices into regions determined to be similar across methods (e.g., the row and column corresponding to a hub node), fitting a convex combination within each partition, and combining these results to yield the ensemble precision matrix.

## Conclusion

The past decade has shown numerous advances in GGM estimation, but the burden has still been left on the researcher to determine the specifics of the estimation process, including important aspects such as choice of method, tuning parameter selection, scoring criteria, and hyperparameter settings. Our SpiderLearner ensemble method removes this barrier, enabling researchers to easily construct a likelihood-based optimal combination from a library of candidate methods. The parsimonious seven-gene risk score identified by our ensemble network-based approach has clear statistical relevance as demonstrated by the validation in six of 15 independent validation datasets, and biological relevance as demonstrated by existing literature on the functions of the seven genes in the SpiderLearner risk score. SpiderLearner is available as open-source R code at https://github.com/katehoffshutta/SpiderLearner, and code to reproduce the simulation and application workflows are available at https://github.com/katehoffshutta/SpiderLearnerWorkflow.

## Supporting information

**S1 Appendix. Asymptotics.** Proof that in the univariate normal case, the negative log likelihood loss function satisfies the bounded tails condition of (25).

**S1 Fig. Comparison of bounded and unbounded loss functions.**

**S2 Fig. Distribution of partial correlations estimated from the CATHGEN metabolomics dataset.**

**S1 Table. Runtimes for a range of** *n,p*.

**S2 Table. Simulation A sensitivity.** Mean sensitivity of edge detection in Simulation A.

**S3 Table. Simulation A specificity.** Mean specificity of edge detection in Simulation A.

**S3 Fig. Simulation A diagnostics.** Relative Frobenius norm, Matrix RV coefficient, in-sample log likelihood, and out-of-sample log likelihood for Simulation A.

**S4 Fig. Simulation A bias and MSE.**

**S4 Table. Simulation B weights.** Average weights of each candidate method in the ensemble for all of the gold-standard networks explored in Simulation B.

**S5 Table. Simulation B sensitivity.** Mean sensitivity of edge detection in Simulation B.

**S6 Table. Simulation B specificity.** Mean specificity of edge detection in Simulation B.

**S5 Fig. Simulation B diagnostics.** Relative Frobenius norm, Matrix RV coefficient, in-sample log likelihood, and out-of-sample log likelihood for Simulation B.

**S6 Fig. Simulation B bias and MSE.**

**S7 Table. Simulation C weights.** Average weights of each candidate method in the ensemble for all of the gold-standard networks explored in Simulation C.

**S8 Table. Simulation C sensitivity.** Mean sensitivity of edge detection in Simulation C.

**S9 Table. Simulation C specificity.** Mean specificity of edge detection in Simulation C.

**S7 Fig. Simulation C diagnostics.** Relative Frobenius norm, Matrix RV coefficient, in-sample log likelihood, and out-of-sample log likelihood for Simulation C.

**S8 Fig. Simulation C bias and MSE.**

**S10 Table. Simulation D sensitivity.** Mean sensitivity of edge detection in Simulation D.

**S11 Table. Simulation D specificity.** Mean specificity of edge detection in Simulation D.

**S9 Fig. Simulation D diagnostics.** Relative Frobenius norm, Matrix RV coefficient, in-sample log likelihood, and out-of-sample log likelihood for Simulation D.

**S10 Fig. Simulation D bias and MSE.**

**S11 Fig. Choice of *K* and variability.** Element-wise standard error as a function of the number of folds *K* used to train the SpiderLearner.

**S12 Fig. Choice of *K* and bias.** Element-wise bias as a function of the number of folds *K* used to train the SpiderLearner.

**S13 Fig. Choice of *K* and variability of ensemble weights.** Boxplots of ensemble weights for each of the nine candidate methods as a function of the number of folds *K* used to train the SpiderLearner.

**S14 Fig. Sensitivity of estimated model to library.** Simulation setting comparing three libraries.

**S15 Fig. Sensitivity of estimated model to candidate method.** Example for easy-to-visualize 14-gene set.

**S12 Table. Sensitivity of estimated model to library.** Table of results from application setting comparing four libraries on small ovarian cancer dataset.

**S16 Fig. Sensitivity of estimated model to library.** Plot of estimated models in application setting comparing four libraries on small ovarian cancer dataset.

**S17 Fig. Ensemble risk score performance in all 15 validation datasets.** Kaplan-Meier plots of high-risk and low-risk groups with log-rank *p*-value.

**S18 Fig. Forest plots of unadjusted and adjusted hazard ratios associated with the SpiderLearner risk score**.

**S19 Fig. Significance of hazard ratios associated with the network risk score for all candidate methods, the simple mean, and the SpiderLearner**.

## Acknowledgments

The authors gratefully acknowledge Subhajit Naskar for his contributions in designing the simulation studies that inspired this work.

The authors gratefully acknowledge the participants of the CATHGEN study and of the 16 ovarian cancer studies utilized in the biological application of this manuscript.

## Author Contributions

KHS conceptualized, developed, and implemented the method. LBB consulted on theoretical foundations of the method. KHS and RB conceptualized the simulation and application. KHS conducted the simulation and application. DMS provided input on the application and interpretation. RB provided guidance and supervision for the project. KHS wrote the manuscript. LBB, DMS, and RB reviewed the manuscript and provided critical input.

## Human Subjects Statement

### CATHGEN dataset

The CATHGEN study was approved by the Duke Institutional Review Board and subjects provided informed consent, as described in (37).

### Ovarian cancer datasets

We refer the reader to each originally published dataset (Table 3) for details regarding human subjects protections in each of these studies. Because all of these data are public, analyses performed here is IRB-exempt under Category 4 - Secondary Research Uses of Identifiable Private Information or Identifiable Biospecimens.

