## Supplement for "SpiderLearner: An ensemble approach to Gaussian graphical model estimation"

July 13, 2021

#### Contents

**1 S1 Appendix: Asymptotics** **1**

**2 Bounded Loss Function Results** **4**

**3 CATHGEN weights** **5**

**4 Simulation Study** **6**

**5 Choice of K** **23**

**6 Selection of Candidate GGM Estimation Methods** **25**

**7 Ovarian Cancer Example** **25**

### 1 S1 Appendix: Asymptotics

The results of (1) require a bounded loss function; however, (2) note that they also hold for certain types of unbounded loss functions as described in (3). Our loss function (negative log likelihood) is clearly unbounded; we therefore need to show that it is one of these types in order to conclude that the oracle results of (1) hold for the SpiderLearner. The specific condition described in (3) is as follows. For a function-specific constant $M(f)$ and every $t > 0$, we must have:

$$P(x : |f(x)| > t) \leq Ce^{-t^p/M(f)^p} \tag{1}$$

for some $p \in [1, 2]$ and where $C$ is a constant (3). This condition is referred to as "exponentially decreasing tails of order $p$". The intuition is that, although the loss function is not bounded, the probability that it will be larger than a certain value can be bounded as in (1).

Using foundational statistical properties, Condition (1) can be easily verified in the case of the log likelihood loss function for a centered normal univariate random variable. Here, we proceed according to the approach demonstrated for sub-Gaussian random variables in (4). Let $X \sim N(0, \sigma^2)$. In this case, the loss function for a single observation $x$ and an estimator $\hat{\sigma}^2$ is:

$$\begin{aligned} Loss(x, \hat{\sigma}^2) &= -\log\left\{\frac{1}{\sqrt{2\pi}}\exp\left(-\frac{x^2}{2\hat{\sigma}^2}\right)\right\} && (2) \\ &= -\log\{\phi(x/\hat{\sigma})\} && (3) \end{aligned}$$

where $\phi(\cdot)$ is the standard Normal density function. We can then write:

$$\begin{aligned} P(x : Loss(x, \hat{\sigma}^2) > t) &= P(-\log\{\phi(x/\hat{\sigma})\} > t) && (4) \\ &= P(\log\{\phi(x/\hat{\sigma})\} < -t) && (5) \\ &= P(\phi(x/\hat{\sigma}) < \exp(-t)) && (6) \\ & && (7) \end{aligned}$$

For convenience, define $a = \phi^{-1}(\exp(-t))$ with $a \geq 0$ (i.e. choose the positive inverse), and let $Z = X/\sigma$. Then we are simply looking for the probability that $|Z| > a$. That probability can be bounded as follows (4; 5):

$$\begin{aligned} P(|Z| > a) &= 2P(Z > a) && (8) \\ &= 2P(\exp(Z) > \exp(a)) && (9) \\ &\leq 2E[\exp(Zs)]/\exp(as) \quad \forall s > 0 \text{ (Markov's inequality)} && (10) \\ &= 2\exp(s^2/2)/\exp(as) \text{ (MGF of Normal distribution)} && (11) \\ &= 2\exp(s^2/2 - as) && (12) \end{aligned}$$

Because this relationship is true for any $s > 0$, we can find the tightest bound by minimizing with respect to $s$ (4):

$$\frac{d}{ds}(\exp(s^2/2 - as)) = \exp(s^2/2 - as) * (s - a) = 0 \text{ when } s = a \qquad (13)$$

Setting $s = a$ yields the bound

$$P(|Z| > a) \leq 2 * \exp(a^2/2 - a^2) = 2\exp(-a^2/2) \qquad (14)$$

With some algebra, we can show that $\phi^{-1}(u) = \pm\sigma\sqrt{-\log(2\pi u^2)}$. Then we can plug in the actual value of $a$:

$$
\begin{aligned}
a &= \sqrt{-\log(2\pi(\exp(-t)^2))} = \sqrt{2t - \log(2\pi)} && (15)\\
P(|Z| > a) &\leq 2\exp\{-(\sqrt{2t - \log(2\pi)})^2/2\} && (16)\\
&= 2\exp(-2t/2 + \log(2\pi)/2) && (17)\\
&= 2\sqrt{2\pi}\exp(-t) && (18)
\end{aligned}
$$

This is of the form in Condition 1, with $C = 2\sqrt{2\pi}$, $p = 1$, and $M(f) = 1$.

In the case of a diagonal covariance matrix, this result extends naturally to the multivariate normal case by factorization of the joint probability density. Our specific interest, however, is in the case of correlated variables. Tail bounds for the multivariate normal density with a non-diagonal covariance matrix are not as straightforward. With the use of Mills's Ratio, a bound can be established under strict conditions on the covariance matrix $\Sigma$ (6). A covariance matrix including both positive and negative entries violates these conditions, and the results of (6) cannot be applied. Some more general bounds have been developed using quadratic programming, but these bounds require the solution of a different quadratic programming problem for every vector-valued $\mathbf{t}$ and do not admit a statement for general $\mathbf{t}$ (7; 8). To our knowledge, no such general statement yet exists. Consequently we cannot apply the asymptotic results of (3) and (1) to the SpiderLearner. We note this as an area for future work and, as discussed in the main paper, suggest the inverse-logit transformation as an alternative for use in practice.

### 2 Bounded Loss Function Results

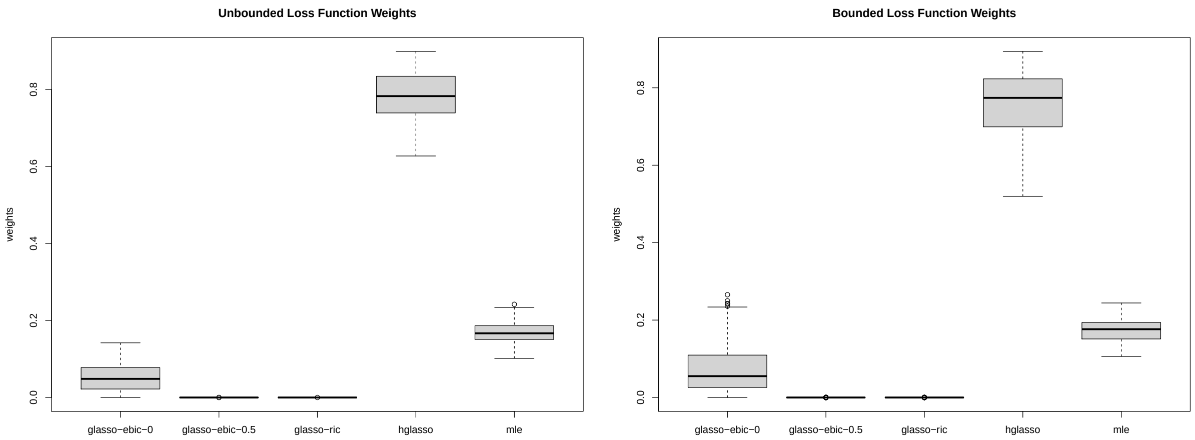

Figure 1: In $n = 100$ simulations with a library of 5 candidates including huge-ebic-0, huge-ebic-0.5, huge-ric, hglasso, and the MLE, we observed similar values for the estimated SpiderLearner weights using the unbounded loss function (left) and bounded loss function (right).

### 3 CATHGEN weights

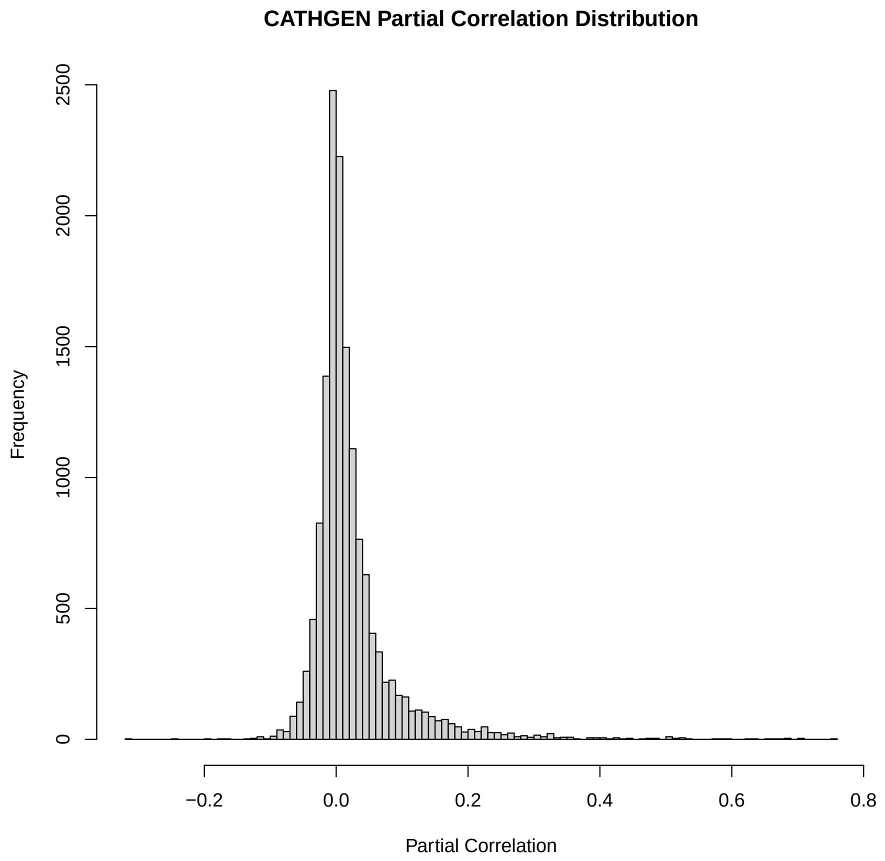

Figure 2: Distribution of nonzero partial correlations estimated from the CATHGEN dataset (9).

### 4 Simulation Study

#### 4.1 Error Metrics

In some cases, it may be of interest to assess the performance of a method at estimating the edge set of the GGM, rather than its edge weights. In this case, GGM estimation is essentially a classification problem, where each possible edge $(i, j)$ is classified as either included in the network or excluded from the network. A typical way of representing the edge set of a network is as a binary adjacency matrix $M$, where $M_{ij} = 1$ if an edge between variable $X_i$ and $X_j$ is included and $M_{ij} = 0$ otherwise. Some methods estimate this matrix directly. Others first estimate a precision matrix, then convert the precision matrix to a matrix of partial correlations in which $M_{ij} \in [0, 1]$, then map these entries to the binary set $\{0, 1\}$ in some way. In Simulations A-C, we follow the latter approach using applying the Fisher transformation to the estimated partial correlation coefficients. The transformation is motivated by the test of the null hypothesis $H_0 : \rho_{ij|X_{-i,-j}} = 0$ vs. the two-sided alternative $H_A : \rho_{ij|X_{-i,-j}} \neq 0$. The Fisher transformation of a sample partial correlation coefficient $\hat{\rho}_{ij|X_{-i,-j}}$ observed for $p-$dimensional data on a sample of size $n$ is given by (10; 11; 12):

$$Z = \frac{1}{2\sqrt{n-3-(p-2)}} \ln\left(\frac{1+\hat{\rho}_{ij|X_{-i,-j}}}{1-\hat{\rho}_{ij|X_{-i,-j}}}\right) \tag{19}$$

Under the null hypothesis of zero partial correlation, the distribution of $Z$ is asymptotically standard normal, facilitating straightforward hypothesis testing. We take advantage of this property by developing the following map to convert the weighted adjacency matrix to a binary adjacency matrix:

$$f(\theta_{ij}) = \begin{cases} 0 & \left|\frac{\theta_{ij}}{\sqrt{\theta_{ii}\theta_{jj})}}\right| < c(\alpha, n, p) \\ \\ 1 & \left|\frac{\theta_{ij}}{\sqrt{\theta_{ii}\theta_{jj})}}\right| \geq c(\alpha, n, p) \end{cases} \tag{20}$$

where $c_\alpha$ is a threshold corresponding to the value of the critical value for a two-sided significance test of the null hypothesis $|\theta_{ij}/\sqrt{\theta_{ii}\theta_{jj})}| = 0$ based on the Fisher-transformed partial correlation (Equation 19). This critical value is based on a test of level $\alpha/(p^2)$ (i.e., a Bonferroni-corrected hypothesis test at level $\alpha$, considering the set of all matrix entries as a collection of multiple tests).

In Simulation D, where $n = 60$ and $p = 100$, $\sqrt{n-3-(p-2)}$ is not a real number. The intuition behind this problem is similar to the reason that ordinary least squares fails when $n < p$, based on the correspondence between partial correlation coefficients and multiple linear regression coefficients (see, e.g., (13).) For this setting, we therefore define edges with the map:

$$f(\theta_{ij}) = \begin{cases} 0 & \theta_{ij} = 0 \\ \\ 1 & \theta_{ij} \neq 0 \end{cases} \tag{21}$$

The comparison of binary adjacency matrices is most naturally viewed in the classification context, where we consider each entry of an estimation matrix as a true positive, false positive, true negative, or false negative. In this way, we can calculate sensitivity and specificity of each estimation method. **Sensitivity** is calculated as the proportion of the true edges that are detected in the estimated adjacency matrix. **Specificity** is calculated as 1 - the proportion of spurious edges detected in the estimated adjacency matrix.

#### 4.2 Runtimes

Runtimes are measured on a MacBook Air with a 1.8 GHz Intel Core i5 processor, and correspond to 10-fold cross-validation.

| n | p | time |
|---|---|---|
| 10000 | 25 | 11.7 min |
| 1000 | 25 | 11.4 min |
| 100 | 25 | 11.3 min |
| 10000 | 75 | 13.3 min |
| 1000 | 75 | 13.6 min |
| 100 | 75 | 15.7 min |
| 10000 | 100 | 15.8 min |
| 1000 | 100 | 14.1 min |
| 100 | 100 | 18.4 min |

Table 1: Runtimes for a range of $n, p$. Runtimes are measured on a MacBook Air with a 1.8 GHz Intel Core i5 processor using only one core, and correspond to 10-fold cross-validation.

##### 4.3 Simulation A

| | ensemble | simple mean | glasso - ebic - 0 | glasso - ebic - 0.5 | glasso - ric | hglasso | mle | glasso - stars - 0.05 | glasso - stars - 0.1 | qgraph - ebic - 0 | qgraph - ebic - 0.5 |
|---|---|---|---|---|---|---|---|---|---|---|---|
| Erdos-Renyi Low | 0.42 | 0.4 | 0.39 | 0.39 | 0.41 | 0.4 | 0.49 | 0.35 | 0.35 | 0.42 | 0.42 |
| Erdos-Renyi High | 0.23 | 0.17 | 0.16 | 0.16 | 0.19 | 0.17 | 0.33 | 0.13 | 0.14 | 0.22 | 0.2 |
| Small World Low | 0.38 | 0.35 | 0.34 | 0.34 | 0.37 | 0.35 | 0.48 | 0.28 | 0.28 | 0.38 | 0.37 |
| Small World High | 0.33 | 0.24 | 0.26 | 0.26 | 0.26 | 0.22 | 0.39 | 0.14 | 0.14 | 0.3 | 0.29 |
| Scale Free Low | 0.31 | 0.29 | 0.3 | 0.29 | 0.29 | 0.28 | 0.41 | 0.23 | 0.25 | 0.3 | 0.29 |
| Scale Free High | 0.26 | 0.19 | 0.2 | 0.2 | 0.2 | 0.18 | 0.34 | 0.13 | 0.14 | 0.23 | 0.21 |
| Hub-and-Spoke Low | 0.36 | 0.32 | 0.31 | 0.31 | 0.35 | 0.33 | 0.45 | 0.28 | 0.28 | 0.39 | 0.37 |
| Hub-and-Spoke High | 0.25 | 0.19 | 0.18 | 0.18 | 0.21 | 0.19 | 0.36 | 0.15 | 0.15 | 0.24 | 0.22 |

Table 2: Mean sensitivity, Simulation A.

| | ensemble | simple mean | glasso - ebic - 0 | glasso - ebic - 0.5 | glasso - ric | hglasso | mle | glasso - stars - 0.05 | glasso - stars - 0.1 | qgraph - ebic - 0 | qgraph - ebic - 0.5 |
|---|---|---|---|---|---|---|---|---|---|---|---|
| Erdos-Renyi Low | 1 | 1 | 1 | 1 | 1 | 1 | 0.9999 | 1 | 1 | 1 | 1 |
| Erdos-Renyi High | 1 | 1 | 1 | 1 | 1 | 1 | 0.9999 | 1 | 1 | 1 | 1 |
| Small World Low | 1 | 1 | 1 | 1 | 1 | 1 | 0.9999 | 1 | 1 | 1 | 1 |
| Small World High | 1 | 1 | 1 | 1 | 1 | 1 | 1 | 1 | 1 | 1 | 1 |
| Scale Free Low | 1 | 1 | 1 | 1 | 1 | 1 | 1 | 1 | 1 | 1 | 1 |
| Scale Free High | 1 | 1 | 1 | 1 | 1 | 1 | 1 | 1 | 1 | 1 | 1 |
| Hub-and-Spoke Low | 1 | 1 | 0.9998 | 0.9998 | 0.9998 | 1 | 1 | 0.9998 | 0.9998 | 1 | 1 |
| Hub-and-Spoke High | 1 | 1 | 1 | 1 | 1 | 1 | 1 | 1 | 1 | 1 | 1 |

Table 3: Mean specificity, Simulation A.

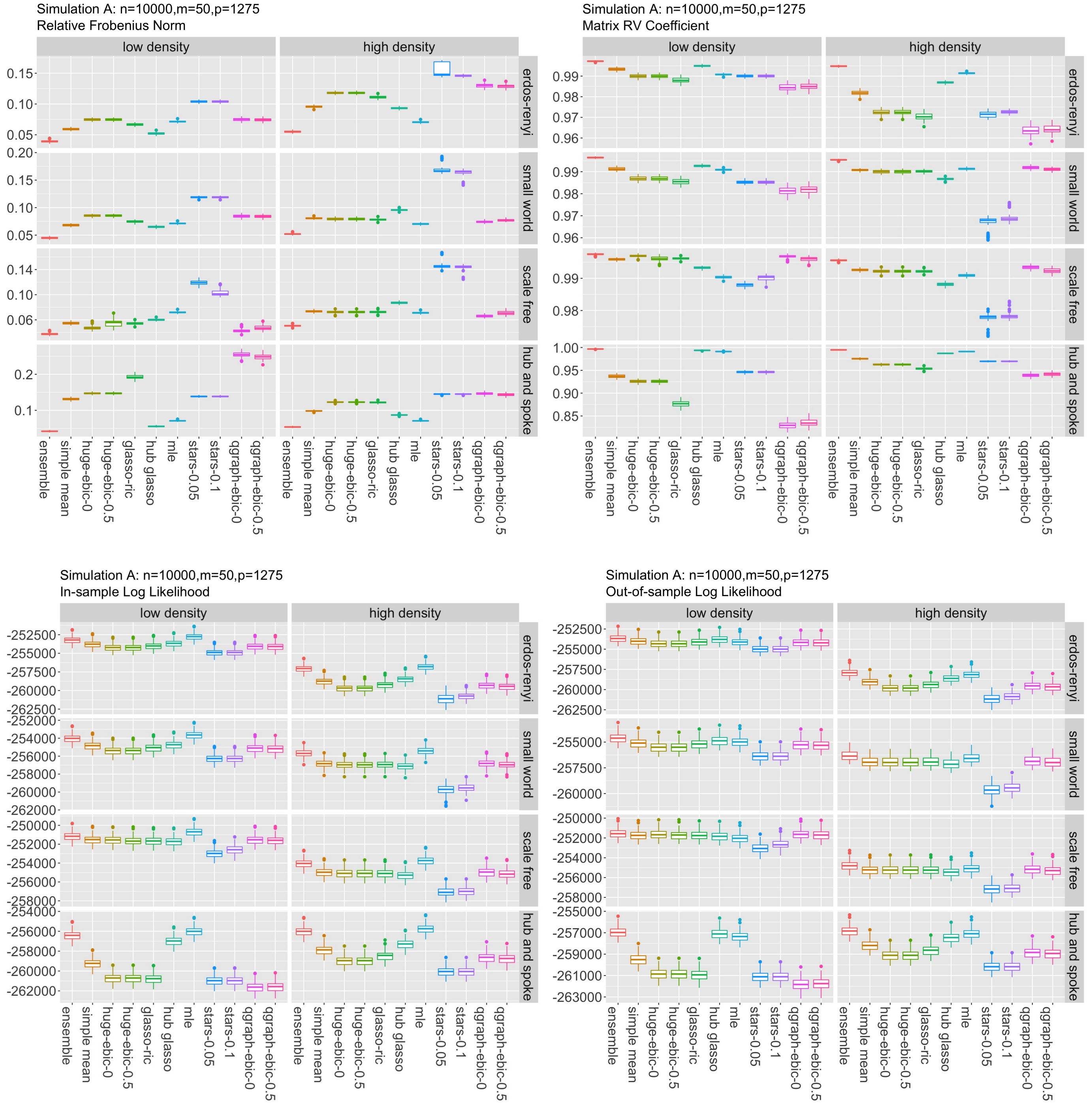

Figure 3: Results for Simulation A.

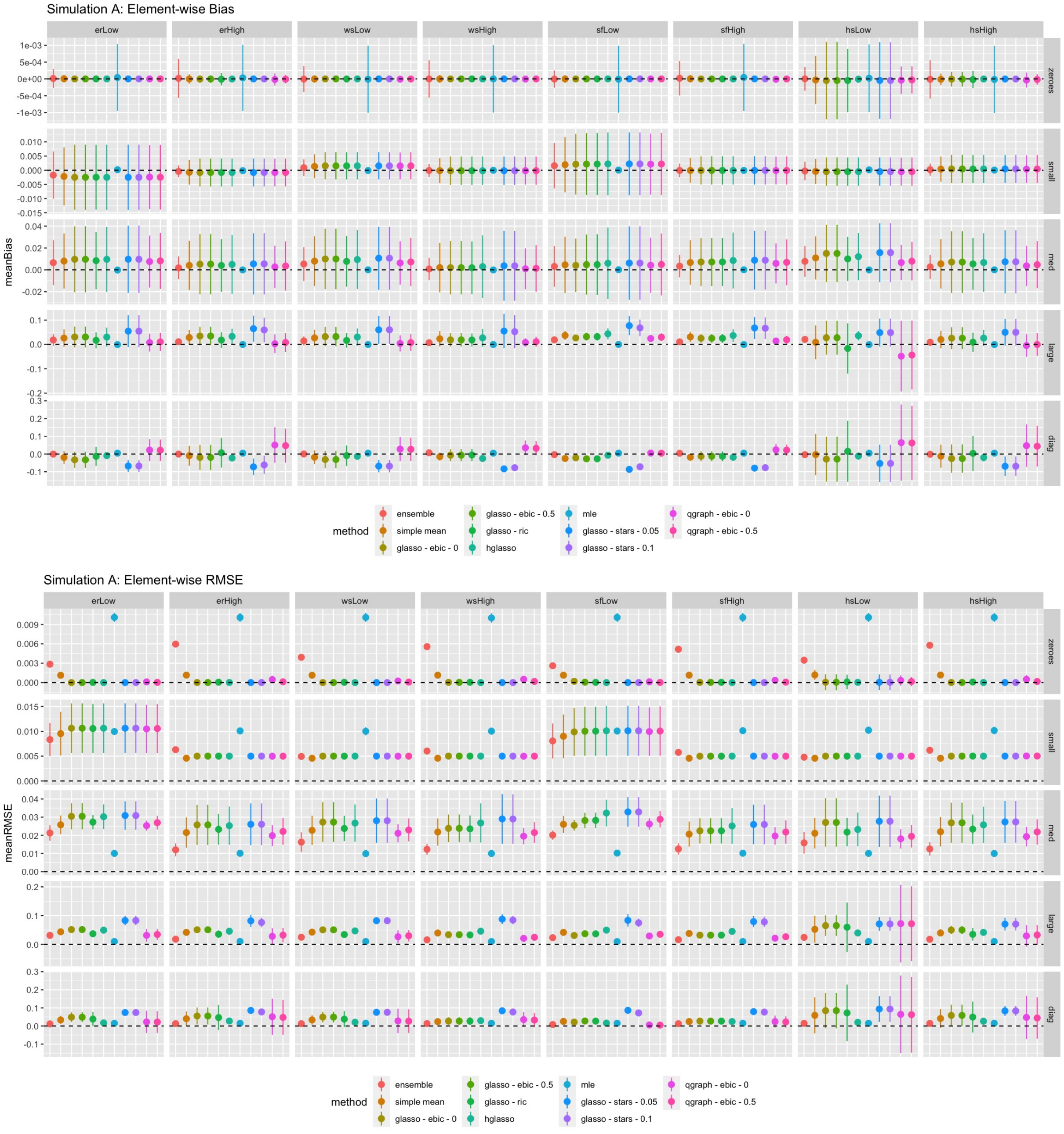

Figure 4: Bias and RMSE for Simulation A.

#### 4.4 Simulation B

| | glasso - ebic - 0 | glasso - ebic - 0.5 | glasso - ric | hglasso | mle | glasso - stars - 0.05 | glasso - stars - 0.1 | qgraph - ebic - 0 | qgraph - ebic - 0.5 |
|---|---|---|---|---|---|---|---|---|---|
| Erdos-Renyi Low | 0.03 | 0 | 0 | 0.76 | 0.03 | 0 | 0 | 0.19 | 0 |
| Erdos-Renyi High | 0.05 | 0 | 0 | 0.79 | 0.14 | 0 | 0 | 0.02 | 0 |
| Small World Low | 0.04 | 0 | 0 | 0.79 | 0.06 | 0 | 0 | 0.11 | 0 |
| Small World High | 0.07 | 0 | 0 | 0.74 | 0.17 | 0 | 0 | 0.02 | 0 |
| Scale Free Low | 0.01 | 0 | 0.05 | 0.46 | 0.06 | 0 | 0 | 0.41 | 0 |
| Scale Free High | 0.08 | 0 | 0 | 0.73 | 0.13 | 0 | 0 | 0.06 | 0 |
| Hub-and-Spoke Low | 0 | 0 | 0 | 0.93 | 0.02 | 0 | 0 | 0.04 | 0 |
| Hub-and-Spoke High | 0.03 | 0 | 0 | 0.82 | 0.12 | 0 | 0 | 0.03 | 0 |

Table 4: Weights for Simulation B

| | ensemble | simple mean | glasso - ebic - 0 | glasso - ebic - 0.5 | glasso - ric | hglasso | mle | glasso - stars - 0.05 | glasso - stars - 0.1 | qgraph - ebic - 0 | qgraph - ebic - 0.5 |
|---|---|---|---|---|---|---|---|---|---|---|---|
| Erdos-Renyi Low | 0.33 | 0.31 | 0.32 | 0.31 | 0.31 | 0.33 | 0.38 | 0.31 | 0.31 | 0.32 | 0.32 |
| Erdos-Renyi High | 0.13 | 0.11 | 0.11 | 0.11 | 0.11 | 0.13 | 0.16 | 0.1 | 0.1 | 0.12 | 0.11 |
| Small World Low | 0.27 | 0.24 | 0.24 | 0.23 | 0.23 | 0.26 | 0.33 | 0.22 | 0.22 | 0.25 | 0.24 |
| Small World High | 0.14 | 0.12 | 0.12 | 0.11 | 0.11 | 0.14 | 0.2 | 0.1 | 0.1 | 0.13 | 0.12 |
| Scale Free Low | 0.22 | 0.21 | 0.21 | 0.21 | 0.21 | 0.23 | 0.28 | 0.2 | 0.21 | 0.21 | 0.21 |
| Scale Free High | 0.13 | 0.12 | 0.12 | 0.12 | 0.12 | 0.13 | 0.17 | 0.1 | 0.11 | 0.12 | 0.12 |
| Hub-and-Spoke Low | 0.27 | 0.25 | 0.26 | 0.25 | 0.25 | 0.27 | 0.31 | 0.24 | 0.24 | 0.26 | 0.26 |
| Hub-and-Spoke High | 0.14 | 0.13 | 0.13 | 0.13 | 0.13 | 0.14 | 0.18 | 0.11 | 0.12 | 0.14 | 0.13 |

Table 5: Mean sensitivity, Simulation B.

| | ensemble | simple mean | glasso - ebic - 0 | glasso - ebic - 0.5 | glasso - ric | hglasso | mle | glasso - stars - 0.05 | glasso - stars - 0.1 | qgraph - ebic - 0 | qgraph - ebic - 0.5 |
|---|---|---|---|---|---|---|---|---|---|---|---|
| Erdos-Renyi Low | 1 | 1 | 1 | 1 | 1 | 1 | 0.9999 | 1 | 1 | 1 | 1 |
| Erdos-Renyi High | 1 | 1 | 1 | 1 | 1 | 1 | 0.9999 | 1 | 1 | 1 | 1 |
| Small World Low | 1 | 1 | 1 | 1 | 1 | 1 | 0.9999 | 1 | 1 | 1 | 1 |
| Small World High | 1 | 1 | 1 | 1 | 1 | 1 | 0.9999 | 1 | 1 | 1 | 1 |
| Scale Free Low | 1 | 1 | 1 | 1 | 1 | 1 | 0.9999 | 1 | 1 | 1 | 1 |
| Scale Free High | 1 | 1 | 1 | 1 | 1 | 1 | 0.9999 | 1 | 1 | 1 | 1 |
| Hub-and-Spoke Low | 1 | 1 | 1 | 1 | 1 | 1 | 0.9999 | 1 | 1 | 1 | 1 |
| Hub-and-Spoke High | 1 | 1 | 1 | 1 | 1 | 1 | 0.9999 | 1 | 1 | 1 | 1 |

Table 6: Mean specificity, Simulation B.

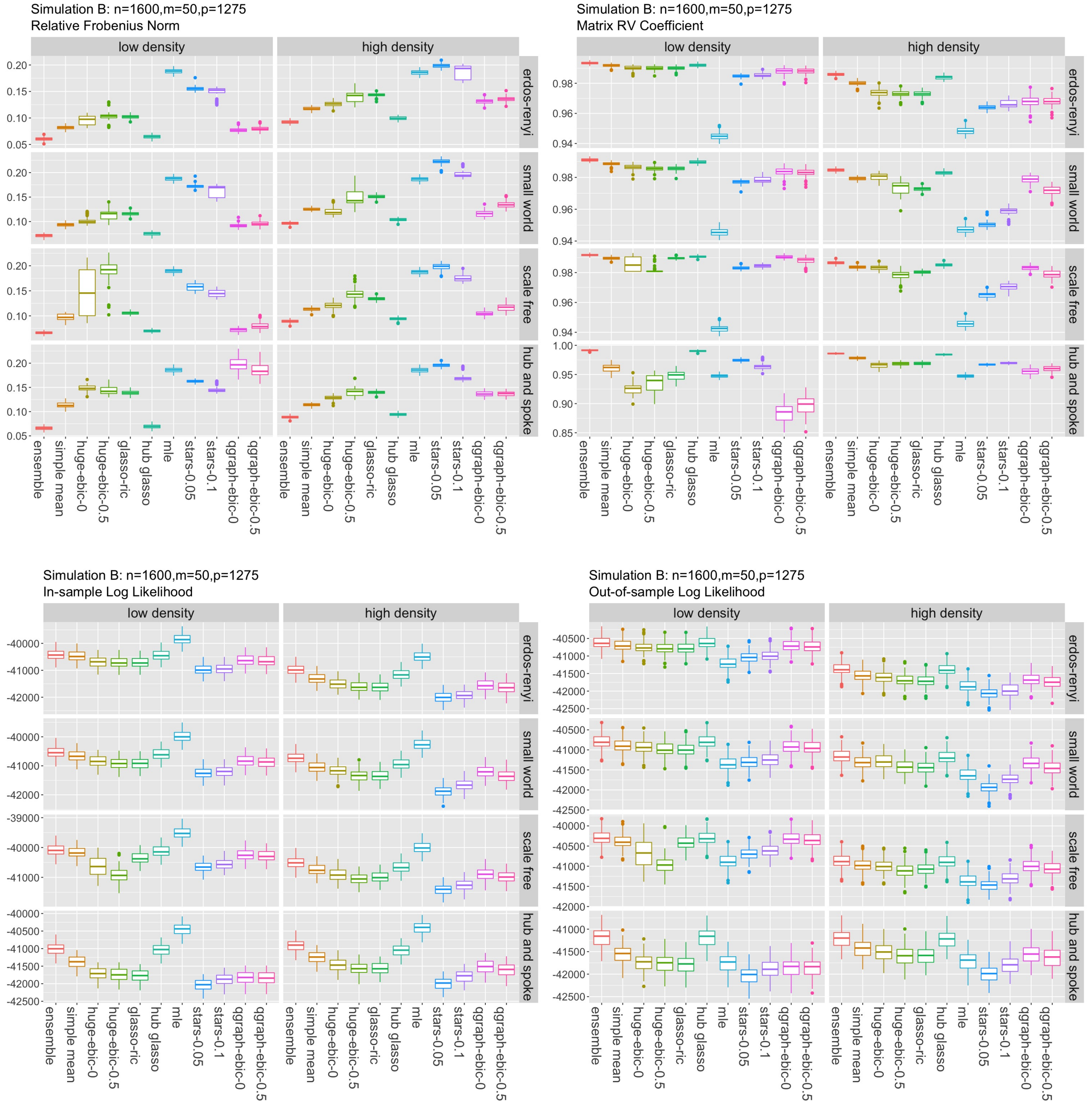

Figure 5: Results for Simulation B.

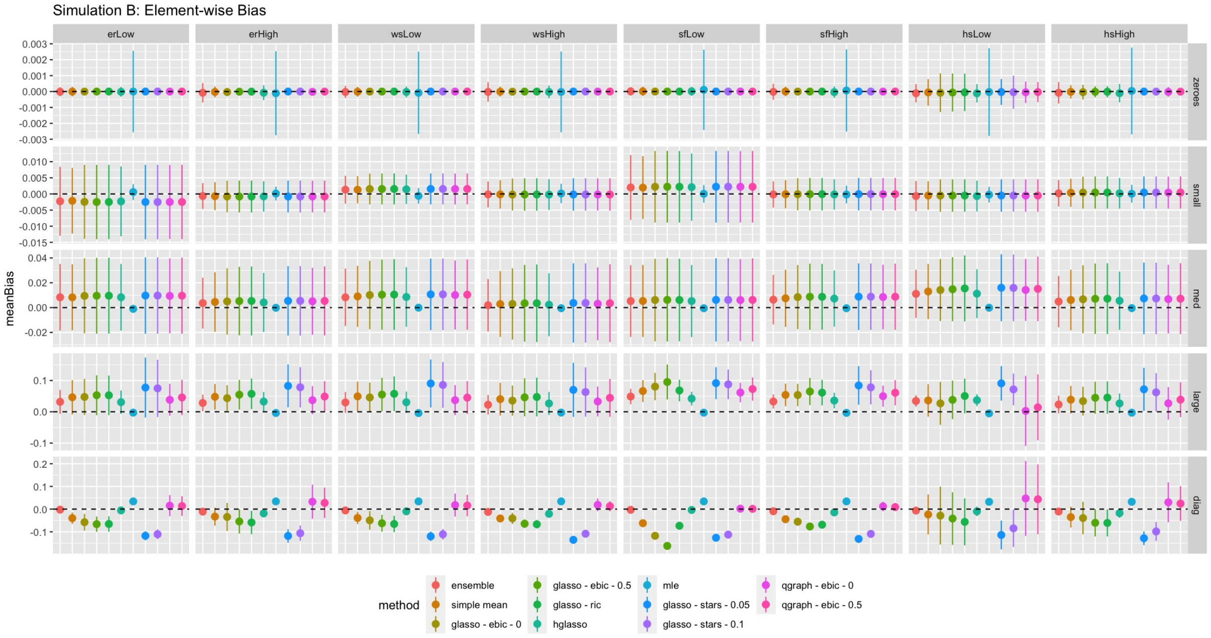

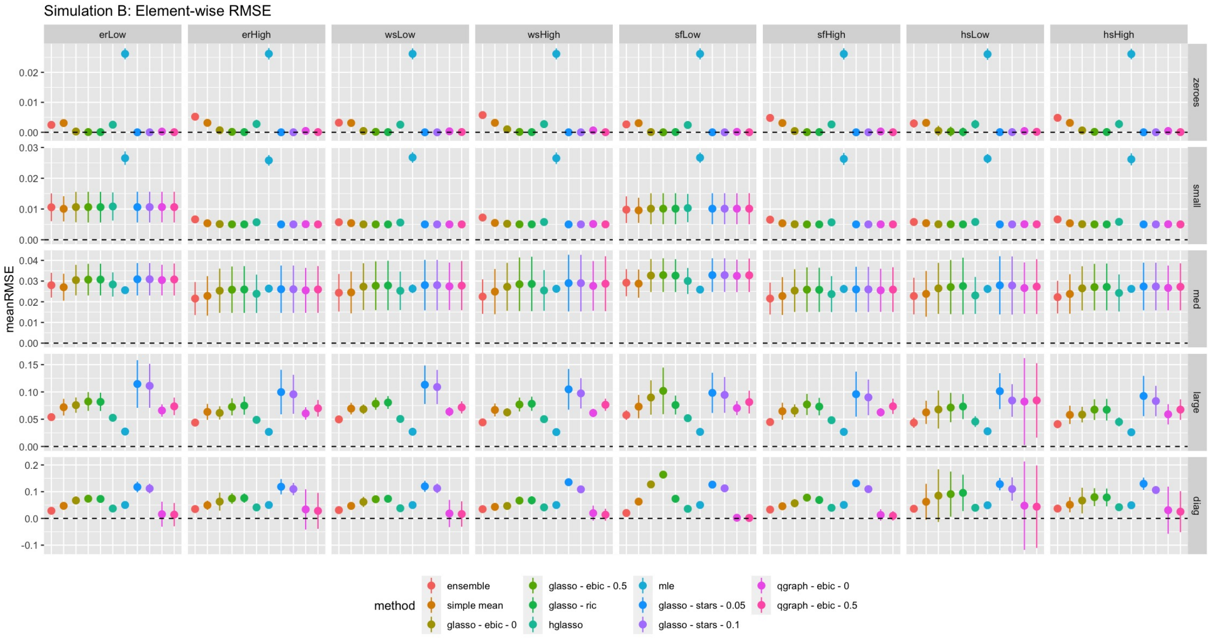

Figure 6: Bias and RMSE for Simulation B.

#### 4.5 Simulation C

| | glasso - ebic - 0 | glasso - ebic - 0.5 | glasso - ric | hglasso | mle | glasso - stars - 0.05 | glasso - stars - 0.1 | qgraph - ebic - 0 | qgraph - ebic - 0.5 |
|---|---|---|---|---|---|---|---|---|---|
| Erdos-Renyi Low | 0 | 0 | 0 | 0.05 | 0 | 0.03 | 0.21 | 0.65 | 0.06 |
| Erdos-Renyi High | 0 | 0 | 0 | 0.1 | 0 | 0.04 | 0.37 | 0.5 | 0 |
| Small World Low | 0 | 0 | 0 | 0.06 | 0 | 0.05 | 0.22 | 0.6 | 0.06 |
| Small World High | 0.01 | 0.01 | 0 | 0.1 | 0 | 0.05 | 0.29 | 0.29 | 0.25 |
| Scale Free Low | 0.02 | 0.02 | 0.01 | 0.03 | 0 | 0.04 | 0.03 | 0.32 | 0.53 |
| Scale Free High | 0.01 | 0.01 | 0 | 0.06 | 0.01 | 0.05 | 0.22 | 0.38 | 0.27 |
| Hub-and-Spoke Low | 0 | 0 | 0 | 0.12 | 0 | 0.02 | 0.39 | 0.46 | 0 |
| Hub-and-Spoke High | 0 | 0 | 0 | 0.11 | 0 | 0.04 | 0.36 | 0.45 | 0.03 |

Table 7: Weights for Simulation C

| | ensemble | simple mean | glasso - ebic - 0 | glasso - ebic - 0.5 | glasso - ric | hglasso | mle | glasso - stars - 0.05 | glasso - stars - 0.1 | qgraph - ebic - 0 | qgraph - ebic - 0.5 |
|---|---|---|---|---|---|---|---|---|---|---|---|
| Erdos-Renyi Low 0.28 | | 0.28 | 0.28 | 0.28 | 0.28 | 0.28 | 0.28 | 0.37 | 0.28 | 0.28 | 0.28 |
| Erdos-Renyi High | 0.09 | 0.09 | 0.09 | 0.09 | 0.09 | 0.09 | 0.18 | 0.09 | 0.09 | 0.09 | 0.09 |
| Small World Low | 0.2 | 0.2 | 0.2 | 0.2 | 0.2 | 0.2 | 0.29 | 0.2 | 0.2 | 0.2 | 0.2 |
| Small World High | 0.09 | 0.09 | 0.09 | 0.09 | 0.09 | 0.09 | 0.18 | 0.09 | 0.09 | 0.09 | 0.09 |
| Scale Free Low | 0.2 | 0.2 | 0.2 | 0.2 | 0.2 | 0.2 | 0.28 | 0.2 | 0.2 | 0.2 | 0.2 |
| Scale Free High | 0.1 | 0.1 | 0.1 | 0.1 | 0.1 | 0.1 | 0.18 | 0.1 | 0.1 | 0.1 | 0.1 |
| Hub-and-Spoke Low | 0.21 | 0.21 | 0.2 | 0.2 | 0.2 | 0.21 | 0.3 | 0.2 | 0.2 | 0.21 | 0.21 |
| Hub-and-Spoke High | 0.1 | 0.1 | 0.1 | 0.1 | 0.1 | 0.1 | 0.19 | 0.1 | 0.1 | 0.1 | 0.1 |

Table 8: Mean sensitivity, Simulation C.

| | ensemble | simple mean | glasso - ebic - 0 | glasso - ebic - 0.5 | glasso - ric | hglasso | mle | glasso - stars - 0.05 | glasso - stars - 0.1 | qgraph - ebic - 0 | qgraph - ebic - 0.5 |
|---|---|---|---|---|---|---|---|---|---|---|---|
| Erdos-Renyi Low | 1 | 1 | 1 | 1 | 1 | 1 | 0.9186 | 1 | 1 | 1 | 1 |
| Erdos-Renyi High | 1 | 1 | 1 | 1 | 1 | 1 | 0.9203 | 1 | 1 | 1 | 1 |
| Small World Low | 1 | 1 | 1 | 1 | 1 | 1 | 0.9206 | 1 | 1 | 1 | 1 |
| Small World High | 1 | 1 | 1 | 1 | 1 | 1 | 0.9222 | 1 | 1 | 1 | 1 |
| Scale Free Low | 1 | 1 | 1 | 1 | 1 | 1 | 0.9223 | 1 | 1 | 1 | 1 |
| Scale Free High | 1 | 1 | 1 | 1 | 1 | 1 | 0.9216 | 1 | 1 | 1 | 1 |
| Hub-and-Spoke Low | 1 | 1 | 1 | 1 | 1 | 1 | 0.9223 | 1 | 1 | 1 | 1 |
| Hub-and-Spoke High | 1 | 1 | 1 | 1 | 1 | 1 | 0.9215 | 1 | 1 | 1 | 1 |

Table 9: Mean specificity, Simulation C.

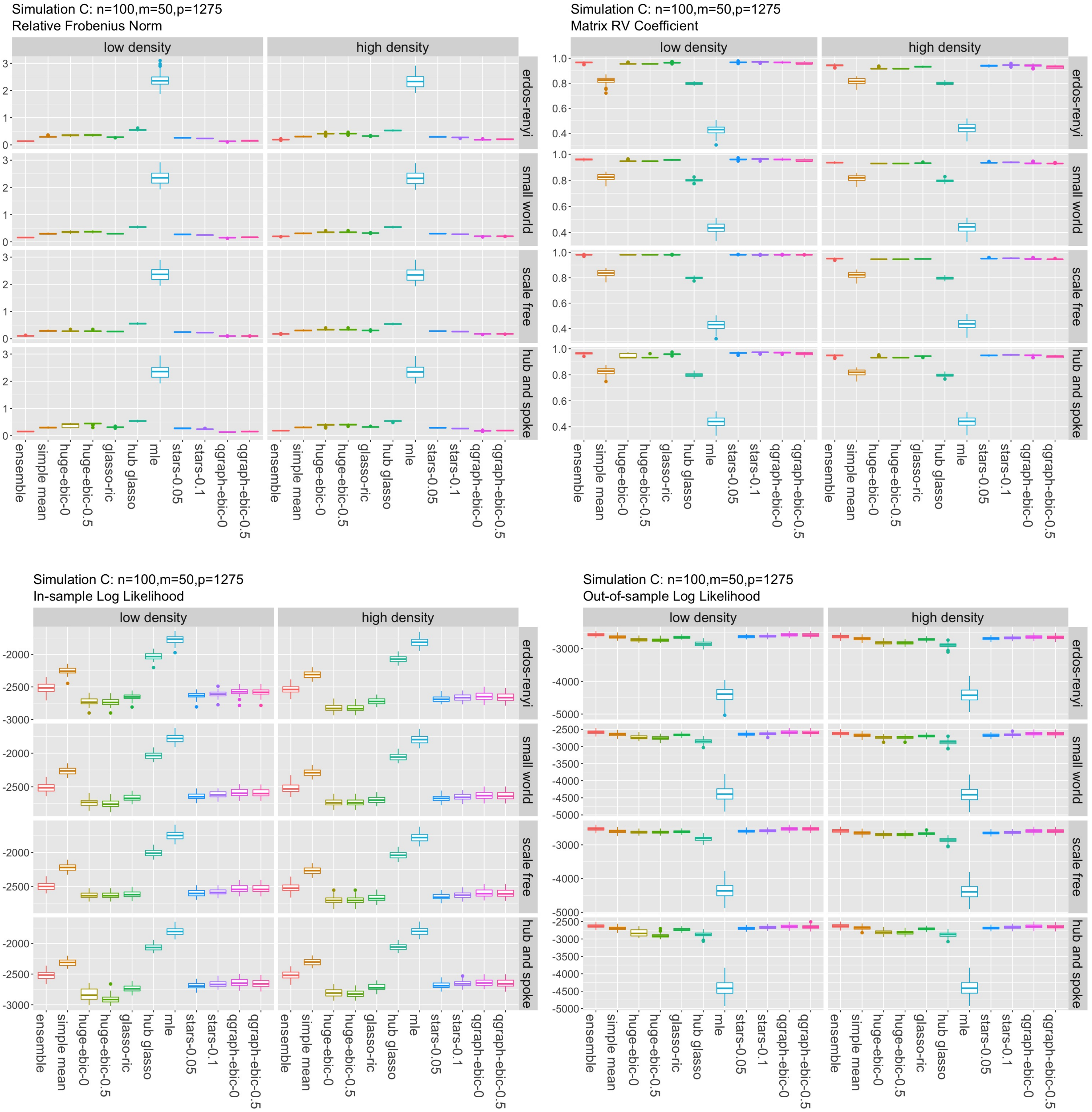

Figure 7: Results for Simulation C.

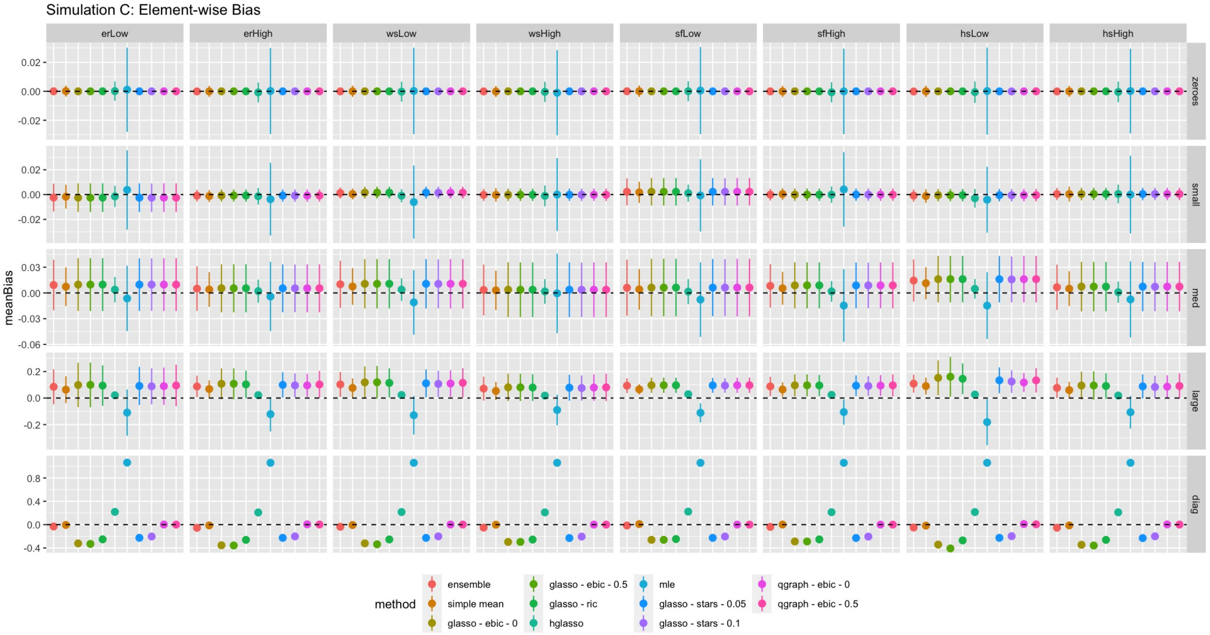

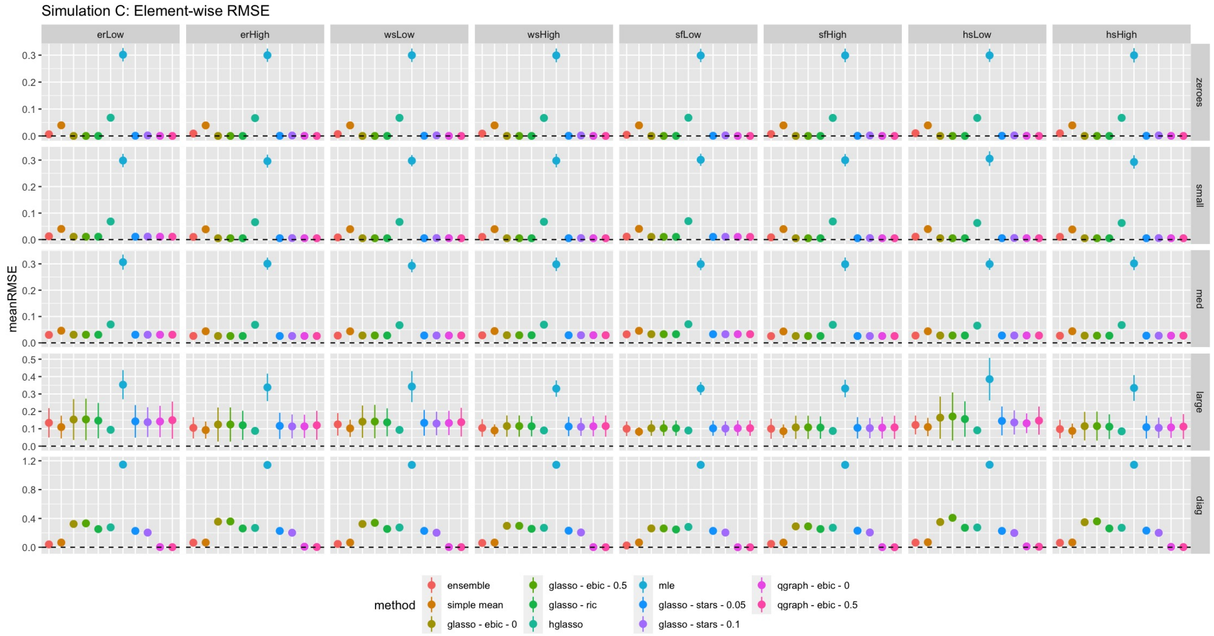

Figure 8: Bias and RMSE for Simulation C.

##### 4.6 Simulation D

| | ensemble | simple mean | glasso - ebic - 0 | glasso - ebic - 0.5 | glasso - ric | hglasso | glasso - stars - 0.05 | glasso - stars - 0.1 | qgraph - ebic - 0 | qgraph - ebic - 0.5 |
|---|---|---|---|---|---|---|---|---|---|---|
| Erdos-Renyi Low | 0.64 | 0.64 | 0.14 | 0.14 | 0.15 | 0.64 | 0.16 | 0.17 | 0.15 | 0.15 |
| Erdos-Renyi High | 0.6 | 0.6 | 0.05 | 0.05 | 0.05 | 0.6 | 0.07 | 0.08 | 0.05 | 0.05 |
| Small World Low | 0.63 | 0.63 | 0.14 | 0.14 | 0.14 | 0.63 | 0.15 | 0.16 | 0.14 | 0.14 |
| Small World High | 0.6 | 0.6 | 0.05 | 0.05 | 0.05 | 0.6 | 0.06 | 0.08 | 0.06 | 0.05 |
| Scale Free Low | 0.63 | 0.63 | 0.15 | 0.15 | 0.15 | 0.63 | 0.16 | 0.17 | 0.15 | 0.15 |
| Scale Free High | 0.61 | 0.61 | 0.05 | 0.05 | 0.05 | 0.61 | 0.08 | 0.1 | 0.07 | 0.05 |
| Hub-and-Spoke Low | 0.64 | 0.64 | 0.15 | 0.15 | 0.15 | 0.64 | 0.16 | 0.17 | 0.15 | 0.15 |
| Hub-and-Spoke High | 0.61 | 0.61 | 0.05 | 0.05 | 0.05 | 0.61 | 0.09 | 0.11 | 0.08 | 0.05 |

Table 10: Mean sensitivity, Simulation D.

| ensemble | simple mean | glasso - ebic - 0 | glasso - ebic - 0.5 | glasso - ric | hglasso | glasso - stars - 0.05 | glasso - stars - 0.1 | qgraph - ebic - 0 | qgraph - ebic - 0.5 | |
|---|---|---|---|---|---|---|---|---|---|---|
| Erdos-Renyi Low | 0.4546 | 0.4546 | 1 | 1 | 1 | 0.4546 | 0.996 | 0.9896 | 0.9998 | 1 |
| Erdos-Renyi High | 0.4484 | 0.4484 | 1 | 1 | 1 | 0.4484 | 0.9964 | 0.9898 | 0.9997 | 1 |
| Small World Low | 0.4573 | 0.4573 | 1 | 1 | 0.9999 | 0.4573 | 0.9964 | 0.9881 | 1 | 1 |
| Small World High | 0.4482 | 0.4482 | 1 | 1 | 1 | 0.4484 | 0.9963 | 0.9854 | 0.9988 | 0.9998 |
| Scale Free Low | 0.456 | 0.456 | 1 | 1 | 1 | 0.456 | 0.9956 | 0.9894 | 0.9999 | 1 |
| Scale Free High | 0.4484 | 0.4484 | 1 | 1 | 1 | 0.4486 | 0.995 | 0.9894 | 0.9983 | 1 |
| Hub-and-Spoke Low | 0.4558 | 0.4558 | 1 | 1 | 0.9999 | 0.4558 | 0.9965 | 0.9879 | 0.9999 | 1 |
| Hub-and-Spoke High | 0.4495 | 0.4495 | 1 | 1 | 1 | 0.45 | 0.9948 | 0.9859 | 0.9976 | 1 |

Table 11: Mean specificity, Simulation D.

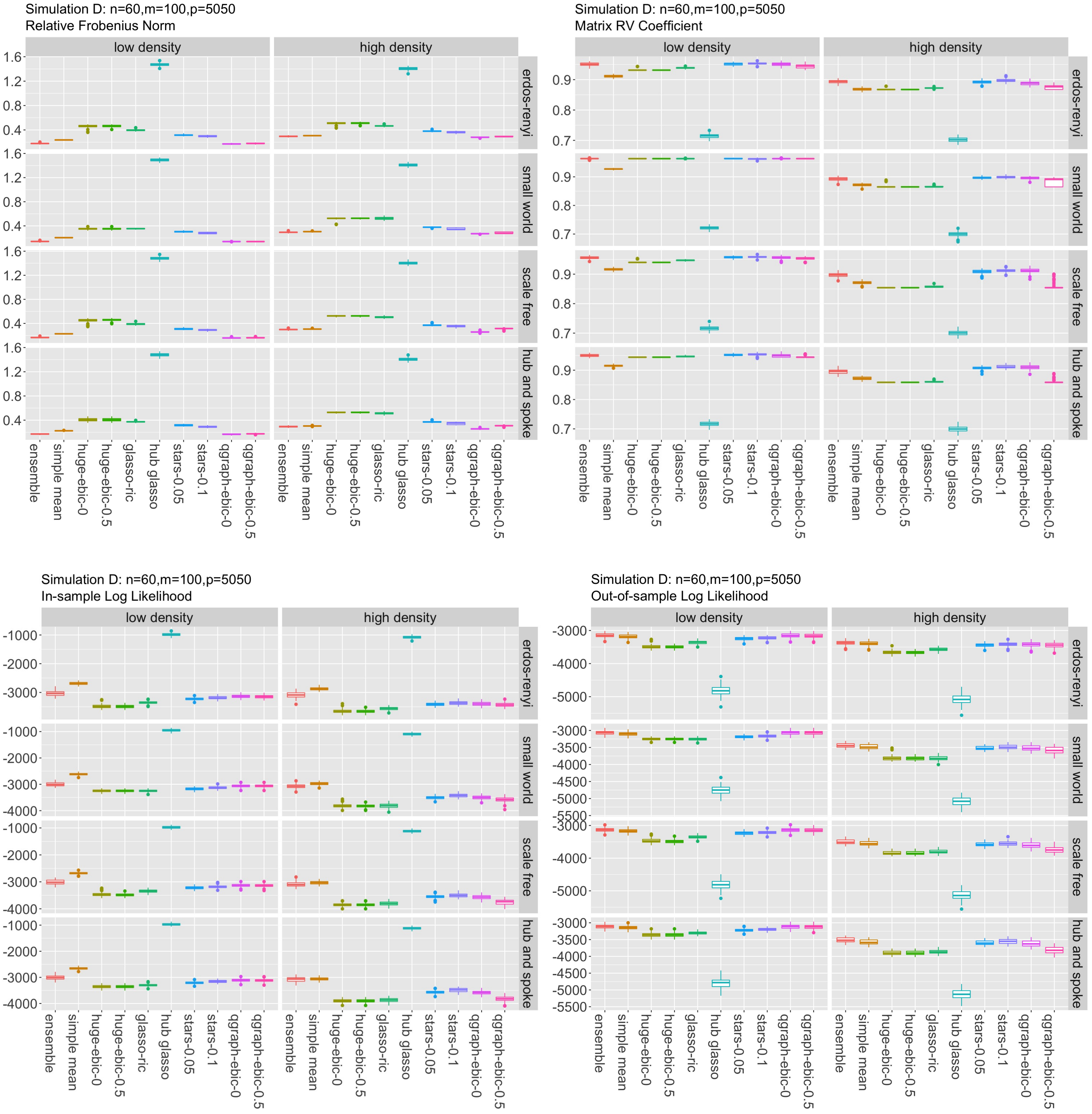

Figure 9: Results for Simulation D.

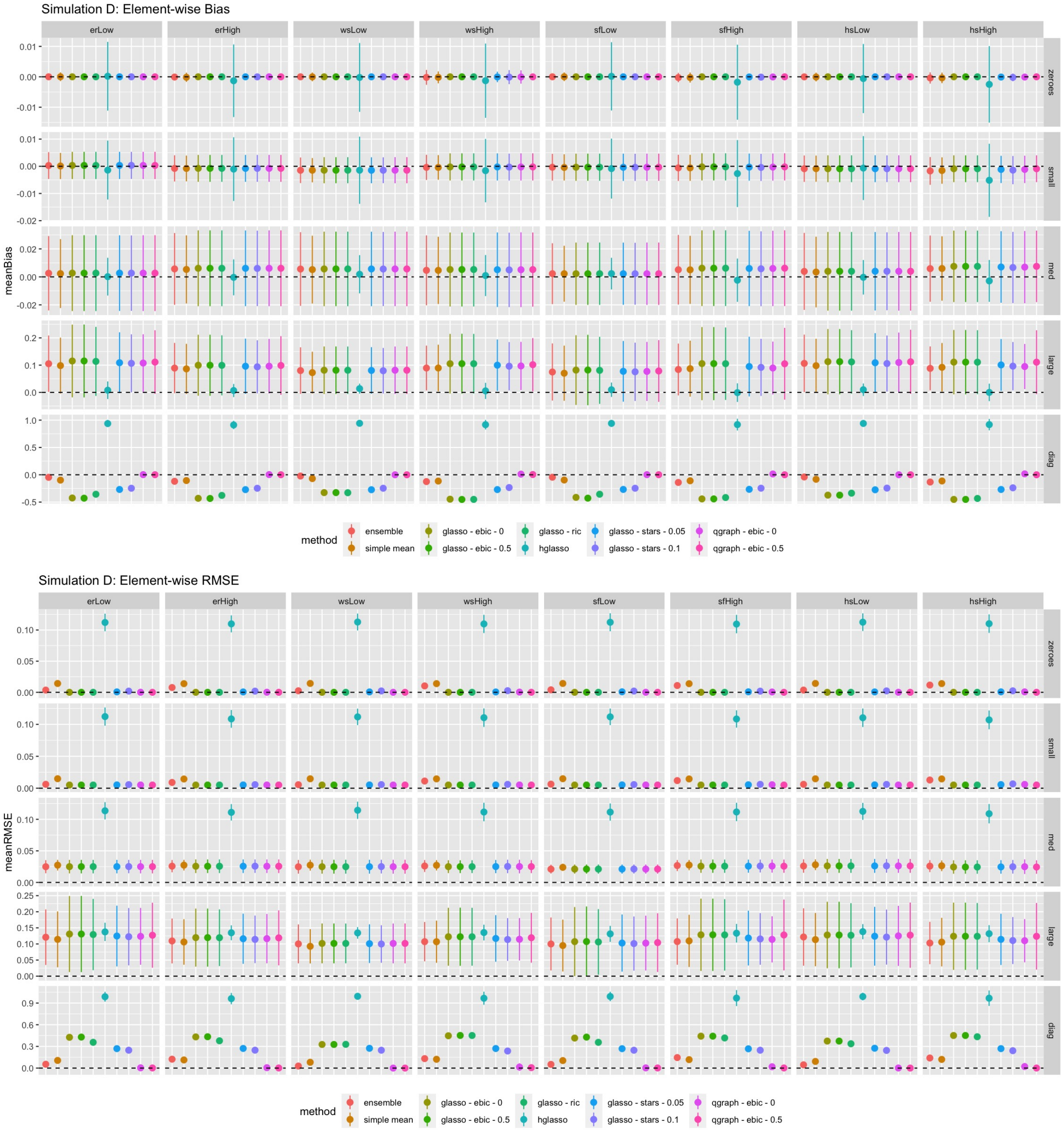

Figure 10: Bias and RMSE for Simulation D.

### 5 Choice of K

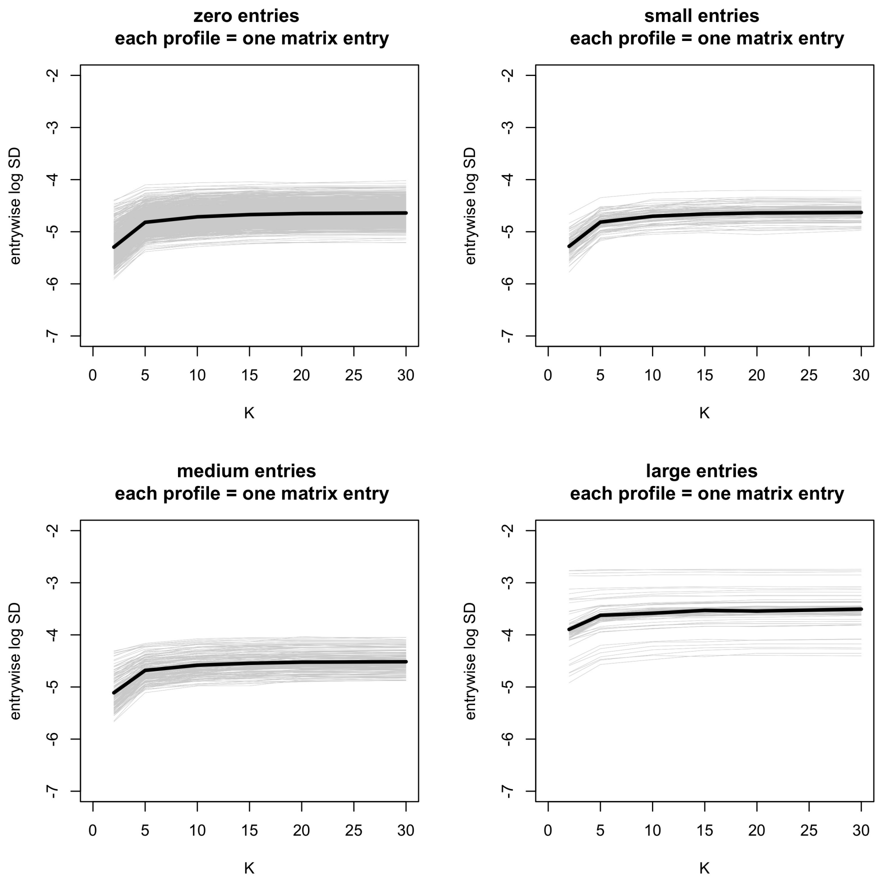

Figure 11: Element-wise standard error as a function of $K$.

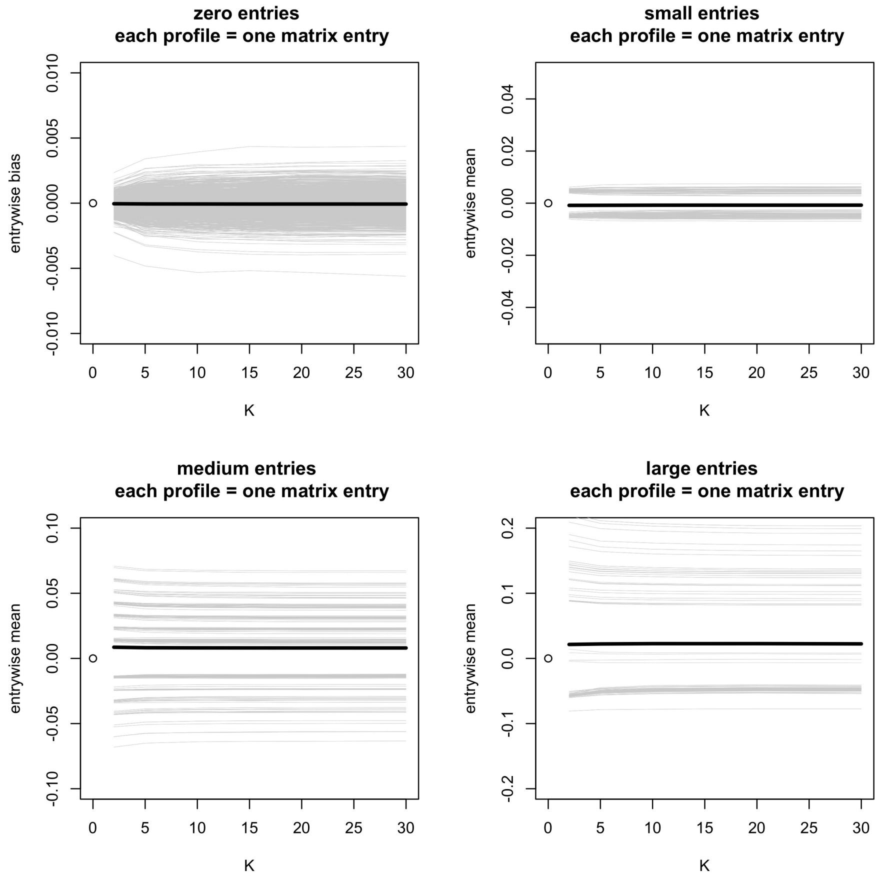

Figure 12: Element-wise bias as a function of $K$.

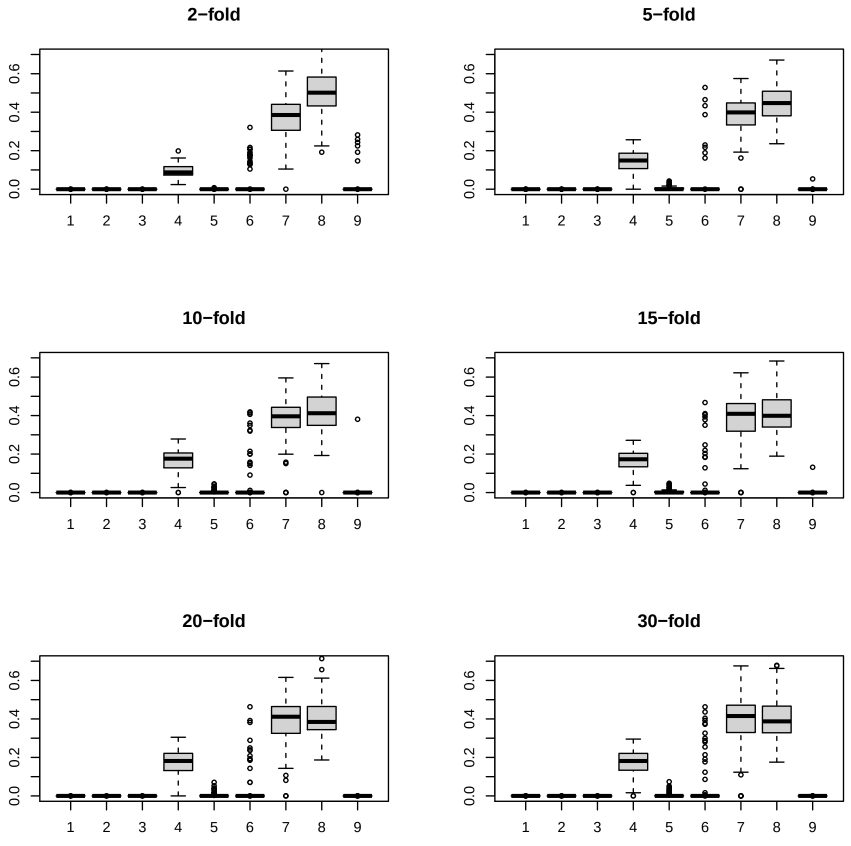

Figure 13: Distribution of weights as a function of $K$. Horizontal axis: nine different models. Vertical axis: estimated weights.

### 6 Selection of Candidate GGM Estimation Methods

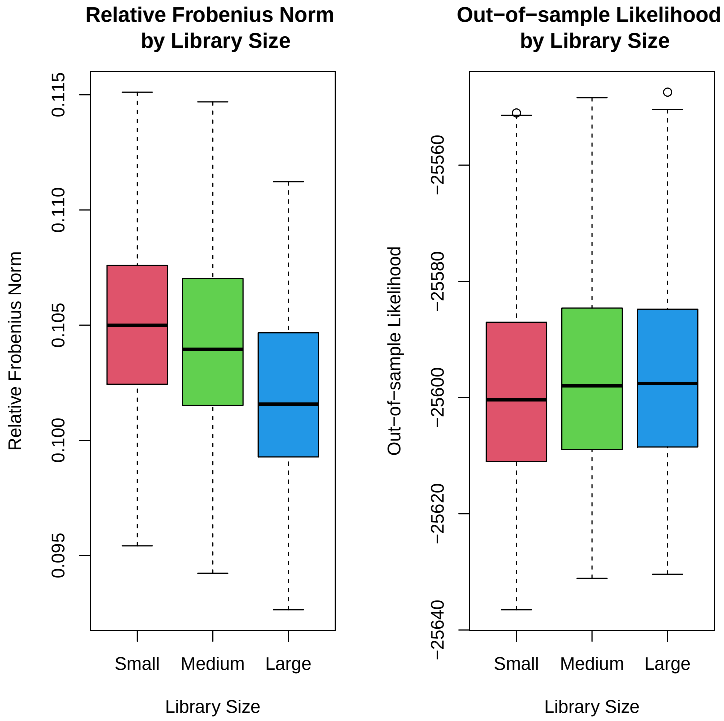

Figure 14: Relative Frobenius norm and out-of-sample log likelihood did not vary substantially with library size in our investigation, although small gains in performance were seen with a larger library.

### 7 Ovarian Cancer Example

#### Ovarian Carcinoma: 14-Gene Set

As a simple example to present additional motivation for the use of an ensemble model like ours, we estimated a GGM on a small subset of the 20106 genes belonging to a gene set corresponding to ovarian carcinoma from the Human Phenotype Ontology (14). This gene set consists of 14 total genes, including 11 tumor suppressor genes (2 of which are also transcription factors), one oncogene, and one protein kinase (15; 16). The SpiderLearner-estimated GGM is shown in Figure 15a, and the seven of the nine candidate models that selected non-empty networks are shown in Figure 15b along with the simple mean model. The estimated GGM is clearly highly sensitive to the choice of estimation method, yet multiple different estimates contributed to the ensemble: the optimal weights for the ensemble model were 0.37 for the `glasso-ebic-0`, 0.26 for the MLE, 0.37 for the `qgraph-ebic-0`, and zero for all the other candidates.

The SpiderLearner model is shown in Supplementary Figure S15a, and the seven of the nine candidate models that selected non-empty networks are shown in Supplementary Figure S15b along with the simple mean model. The estimated GGM is clearly highly sensitive to the choice of estimation method, and multiple different estimates contributed to the optimal ensemble: the weights for the SpiderLearner were 0.37 for the `glasso-ebic-0`, 0.26 for the MLE, 0.37 for the `qgraph-ebic-0`, and zero for all the other candidates.

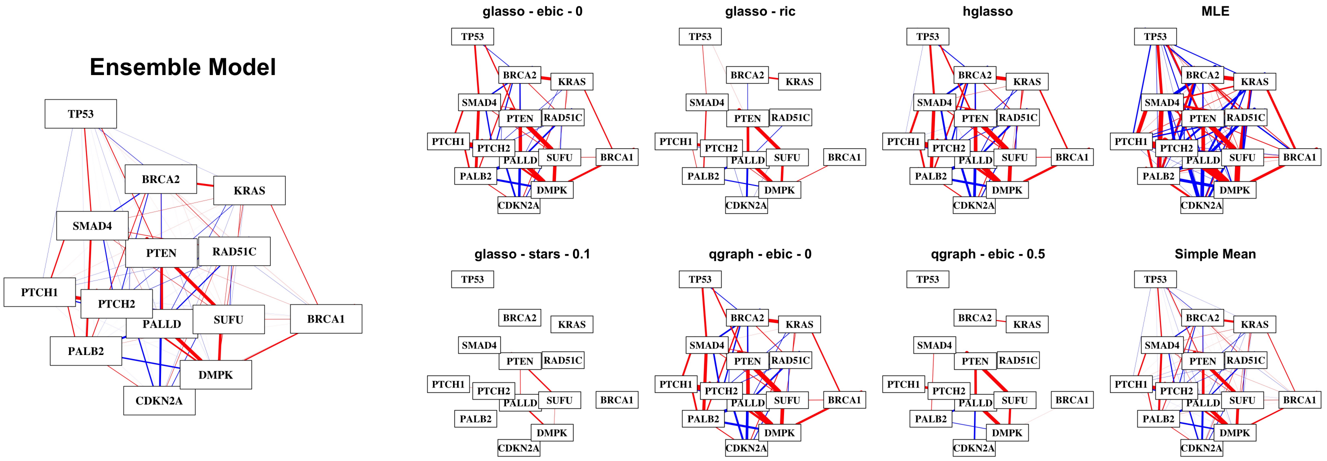

(a) Ensemble model

(b) Candidate models and simple mean

Figure 15: Application of the SpiderLearner method and candidate algorithms to genes contained in an ovarian carcinoma gene set demonstrates the variability of GGM estimation in a practical setting.

**Sensitivity to library selection** To explore the sensitivity of the estimated network to the candidates included in the library, we investigated a variety of library sizes (Supplementary Table S12a). In Model 1, we included all nine candidates used in the simulation study. The optimal weights for these candidates were 0.39 for `huge-ebic-0`, 0.19 for the MLE, 0.42 for the `qgraph-ebic-0`, and zero for the remaining candidates. In Model 2, we considered a "screened" library containing the three candidates with nonzero weights in Model 1. In Model 3, we used a library containing the six candidates that were assigned weights of zero in Model 1. Finally, in Model 4, we removed any candidates with nonzero weight in Model 3, in an effort to create a model without very many strong candidates in the library. With this heavily reduced library, the `qgraph-ebic-0.5` was weighted 1.00 and no other candidates were included in the ensemble.

One way of assessing the differences between two estimated precision matrices is to calculate the relative Frobenius norm between them. Although there is no way to tell which model is best in a real data setting, this calculation permits an understanding of how much an estimate changes with a specific library change. In Table 12b, we show this quantity for pairwise comparisons of the library selections described above. The relative Frobenius norm between Model 1 and Model 2 is near that between Model 2 and Model 3, suggesting that removing the zero-weighted candidates from the library does in fact change the estimate, and the effect size is comparable to that of removing the top-weighted candidates from the library - however, it is a relatively small change when compared to Model 4, which is very different from the other three models according to this measure. A visual comparison of the networks fit by these four models is shown in Supplementary Figure S16.

| Method | Model 1 | Model 2 | Model 3 | Model 4 |
|---|---|---|---|---|
| huge-ebic-0 | 0.39 | 0.79 | x | x |
| huge-ebic-0.5 | 0 | x | 0 | 0 |
| huge-ric | 0 | x | 0.17 | x |
| hglasso | 0 | x | 0.83 | x |
| mle | 0.19 | 0.17 | x | x |
| stars-0.05 | 0 | x | 0 | 0 |
| stars-0.1 | 0 | x | 0 | 0 |
| qgraph-ebic-0 | 0.42 | 0.04 | x | x |
| qgraph-ebic-0.5 | 0 | x | 0 | 1 |

(a) Weights of different methods with a variety of library selections.

| | Model 1 | Model 2 | Model 3 | Model 4 |
|---|---|---|---|---|
| Model 1 | 0.00 | 0.04 | 0.04 | 0.26 |
| Model 2 | 0.04 | 0.00 | 0.03 | 0.22 |
| Model 3 | 0.04 | 0.03 | 0.00 | 0.22 |
| Model 4 | 0.23 | 0.20 | 0.20 | 0.00 |

(b) Relative Frobenius norm calculated between matrices estimated with the libraries described above.

Table 12: Four different library sizes were used to calculate networks on the 14-gene ovarian cancer example. Model 1 contains a full library of 9 candidate algorithms. Model 2 contains only those algorithms that had non-zero weights in Model 1. Model 3 contains only those algorithms that had zero weight in Model 1. Model 4 contains the algorithms that had zero weight in both Model 1 and Model 3. We see robustness in estimation across the first three libraries considered (Model 1, Model 2, Model 3), while the network estimated by Model 4 varies substantially from the first three.

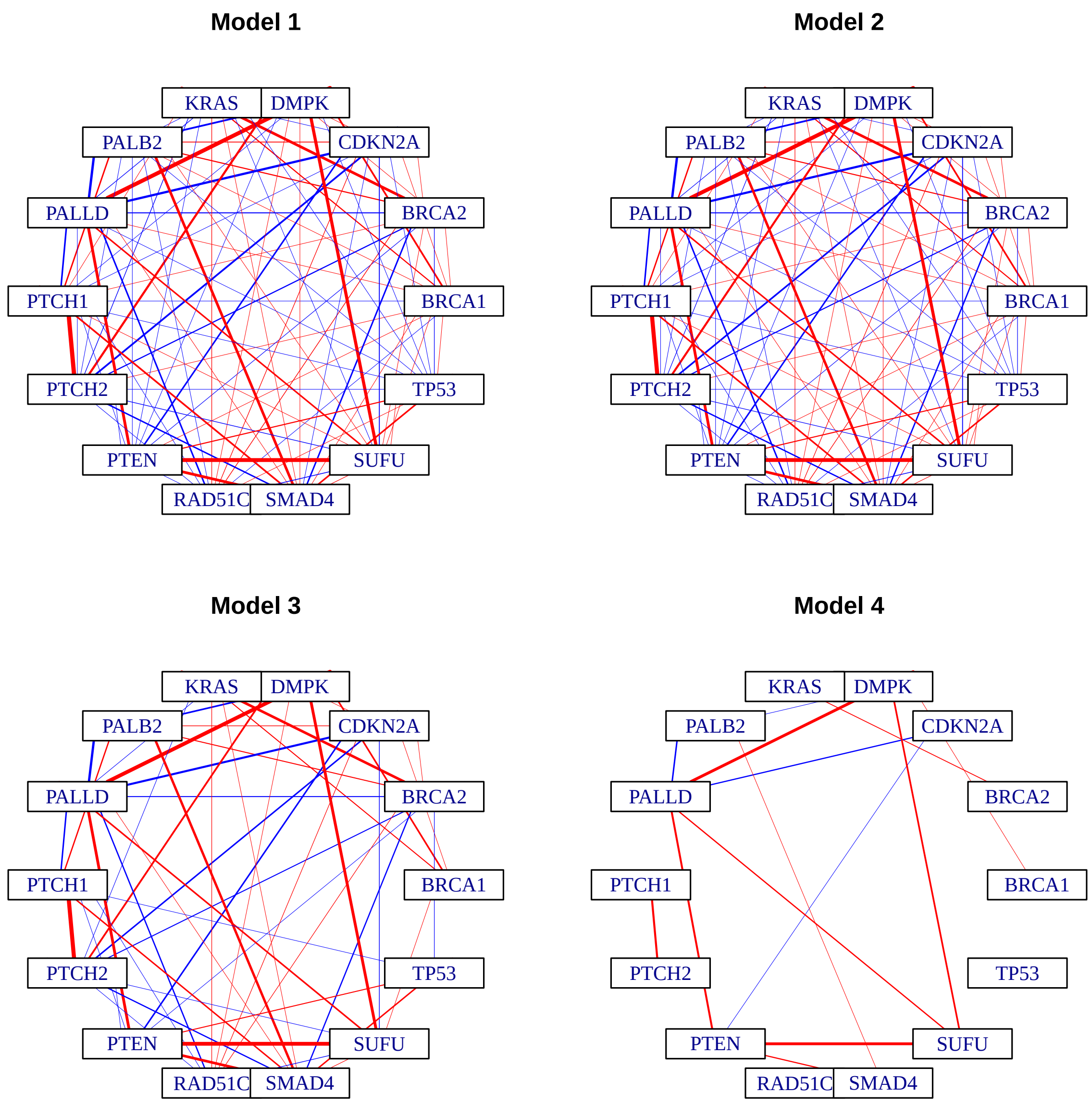

Figure 16: Sensitivity of the estimated network to the library of candidate methods.

Our recommendation, in keeping with (1; 2; 17), is to include as broad of a library of candidates as possible. In our experience, this is not prohibitively computationally expensive. However, if bootstrap-based confidence intervals are desired, computing time can quickly become an issue. In this case, one may wish to use this initial fitting to ”screen” candidates, then perform the bootstrap with only those candidates that had non-zero weights in the first round. On a MacBook Air with a 1.8 GHz Intel Core i5 processor, Model 1 takes 554 seconds to run on two cores while Model 2 takes 19.3 seconds on two cores. If one were to do 100 bootstrap iterations, that time difference would become 15.3 hours vs. 32.1 minutes. We recommend visual and numeric comparison of the networks estimated with the full and screened libraries to assess sensitivity prior to using this screening strategy, and note that because this strategy involves “looking twice” at the data, confidence intervals estimated with this methodology may not have proper coverage. This is an area for future work. An alternative recommendation to this screening approach is to use the parallel processing feature of the SpiderLearner code and run the process on multiple cores.

### High-Risk Ovarian Cancer Signature: 126-Gene Set

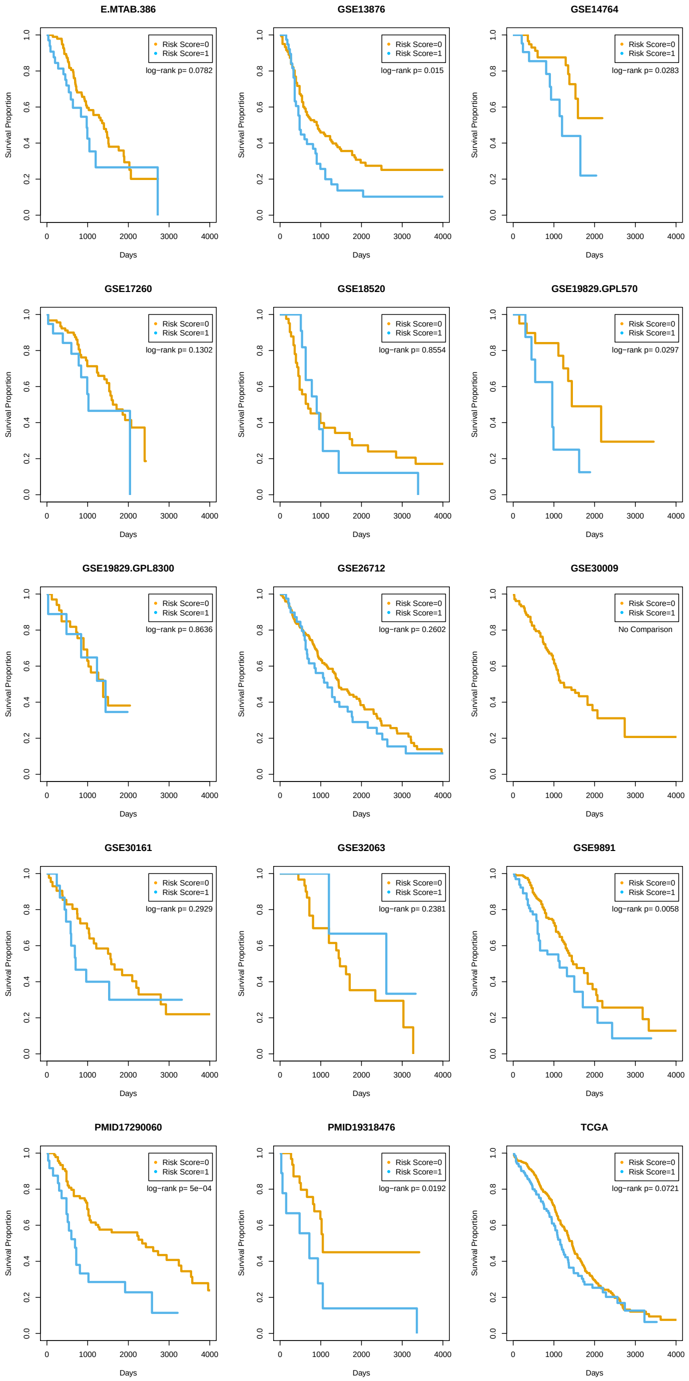

Figure 17: Kaplan-Meier curves for all 15 validation datasets using the ensemble-trained risk score.

**Cox PH Model Results**

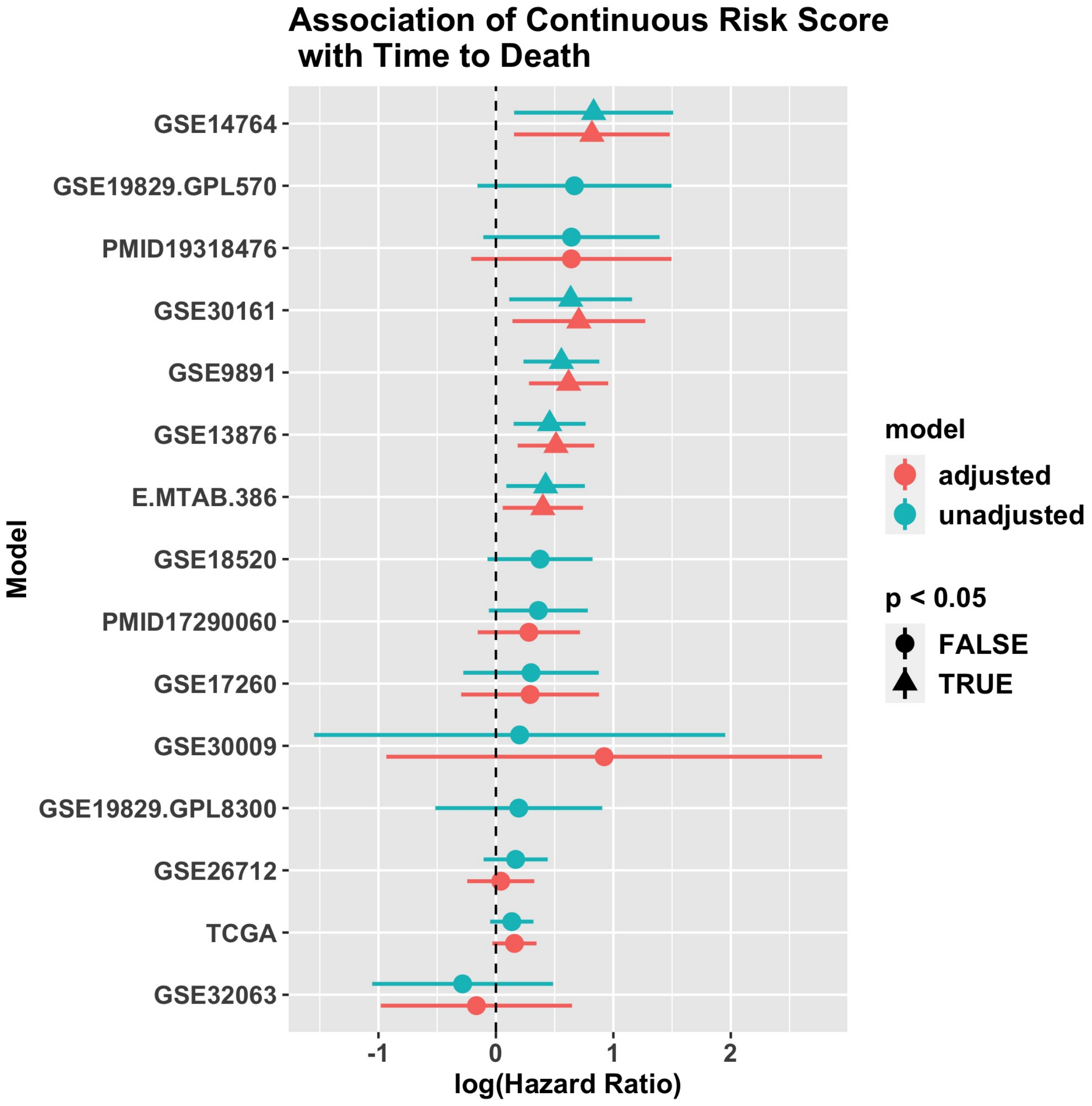

Figure 18: Unadjusted and adjusted log hazard ratios for the SpiderLearner risk score in Cox PH models. Covariates used for adjustment can be found in Table 5 of the main manuscript.

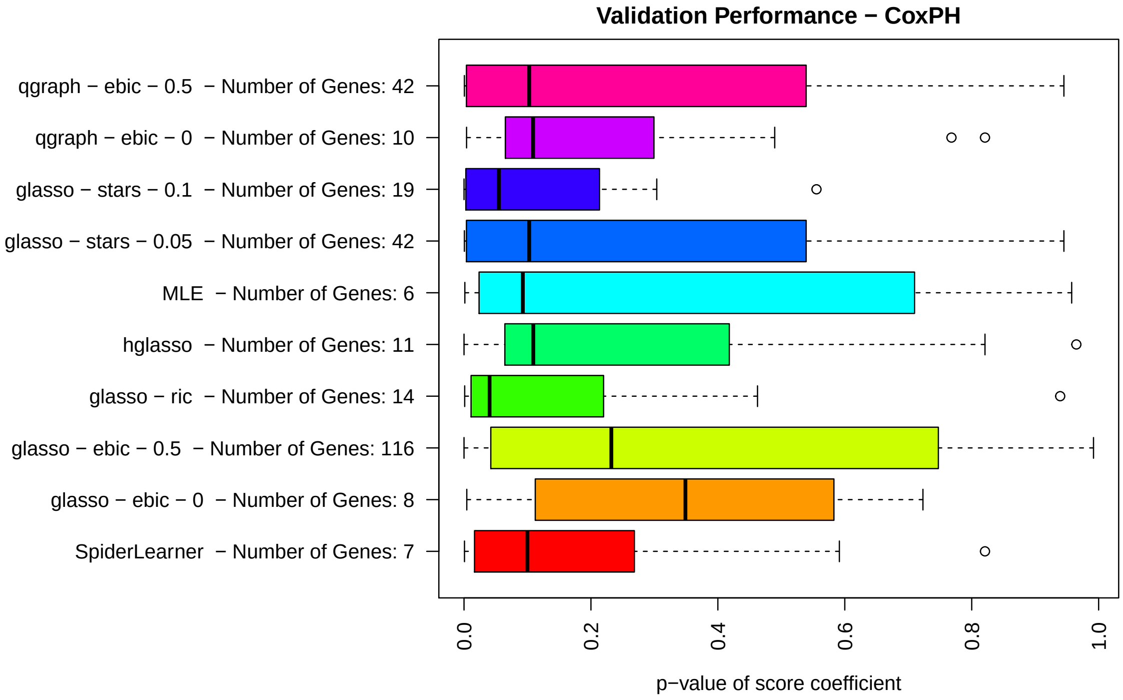

Figure 19: Distribution of p-values for the coefficient of the risk score developed with each method in unadjusted Cox PH models.
